# Prior Context Scaffolds Sentential Semantic Integration during Noisy Speech Comprehension

**DOI:** 10.64898/2026.07.11.737996

**Authors:** Xinmiao Zhang, Zhuoran Li, Dan Zhang

## Abstract

Understanding speech in noise is a central challenge of everyday communication, yet listeners often succeed by using prior context. How the brain uses such context remains debated: it may refine predictions about upcoming words, or it may provide a higher-level framework that helps degraded speech cohere into meaning. Here we combined simultaneous EEG-fNIRS recording with hierarchical multivariate encoding models to track how prior context shapes speech processing from acoustics to words and sentential meaning. Participants listened to natural spoken narratives under clear speech, noisy speech, and context-supported noisy speech conditions. Context brought comprehension of noisy speech close to clear-speech levels. EEG revealed that contextual support reduced neural encoding of lexical surprisal and entropy, indicating weaker tracking of local word-level prediction demands. In contrast, when context was available, fNIRS showed enhanced encoding of sentence-level semantic integration across frontal regions and the right angular gyrus, and stronger angular gyrus encoding predicted better comprehension. By combining EEG and fNIRS to capture complementary electrophysiological and hemodynamic signals, this multimodal approach reveals a hierarchical shift in degraded speech comprehension: prior context does not simply improve word-by-word prediction, but scaffolds the integration of noisy input into coherent discourse.

## Introduction

Understanding speech in noisy environments is a common challenge of everyday communication. In a crowded room, on a busy street, or through a poor audio connection, the acoustic signal reaching the listener is often incomplete or unreliable. Yet comprehension frequently remains possible because speech is rarely interpreted as an isolated stream of sounds. Listeners can draw on prior knowledge, the topic of conversation, and the broader meaning of the unfolding discourse to infer what is being said. Behavioral and neurocognitive studies have shown that contextual information, including familiar topics, preceding linguistic context, or a brief synopsis, can facilitate the perception and comprehension of degraded speech [1–5]. However, the neural mechanisms through which prior context supports speech-in-noise comprehension remain incompletely understood.

One influential possibility is that prior context helps listeners by strengthening prediction. During language comprehension, the brain continuously uses preceding contextual information to anticipate upcoming input [6–9]. A constraining context can reduce uncertainty about likely words and make unexpected words more salient, thereby supporting interpretation when the acoustic signal is unreliable. This predictive mechanism may be especially useful in noisy listening, where bottom-up sensory evidence is degraded [1,10,11] and listeners may need to rely more strongly on top-down expectations to identify words and resolve ambiguity [12,13]. From this perspective, informative prior context could facilitate comprehension by increasing lexical predictability and reducing uncertainty about upcoming linguistic input.

A second, complementary possibility is that prior context supports comprehension by providing a semantic scaffold. Rather than primarily improving word-by-word prediction, context may offer a higher-level framework that helps listeners organize degraded speech into a coherent representation of the speaker’s intended meaning. Schema-based theories propose that prior knowledge can provide an organizing structure within which new information is interpreted, integrated, and remembered [14–17]. This mechanism may be particularly relevant for naturalistic speech, where comprehension depends not only on identifying individual words but also on integrating sentences over time, maintaining the topic of the discourse, and linking new information to what has already been understood [18,19]. Consistent with this view, previous studies have implicated regions such as the angular gyrus, a region associated with semantic integration, in using semantic context to support speech perception under noisy or degraded conditions [20–22]. Thus, a brief synopsis may help listeners interpret noisy speech by guiding sentence-level and discourse-level meaning construction.

Although both prediction and semantic scaffolding are plausible mechanisms, their relative contributions to contextual facilitation during naturalistic speech-in-noise comprehension remain unclear. Many previous studies have used isolated words, word pairs, or short sentences, which are well suited for testing specific mechanisms but may not fully capture the extended, incremental, and discourse-level processes involved in real-life listening [23,24]. Moreover, prior work has often focused on a single level of processing, such as acoustic tracking, lexical prediction, or semantic integration [20,25,26]. As a result, it remains difficult to determine whether prior context facilitates degraded speech comprehension by broadly enhancing neural encoding across processing levels, strengthening local lexical prediction, or primarily supporting higher-level semantic integration.

Addressing this question requires both naturalistic materials and a framework that can separate neural responses to different levels of speech information. Multivariate temporal response function modeling provides such an approach by estimating how neural activity tracks continuous stimulus features over time [27,28]. This framework has been widely used to examine neural encoding of acoustic, lexical, and semantic features during continuous speech comprehension [29–32]. In the present study, we used this approach to dissociate neural encoding of acoustic, lexical, and sentential features during narrative listening. Acoustic envelope encoding indexed tracking of low-level speech fluctuations; lexical surprisal and entropy indexed word-level prediction error and uncertainty; and sentence-level semantic dissimilarity indexed how each incoming sentence was integrated with the evolving discourse context.

We further combined this hierarchical encoding approach with simultaneous EEG-fNIRS recording. EEG captures electrophysiological responses with high temporal sensitivity, making it well suited for tracking rapid acoustic and lexical dynamics during continuous speech [29,30]. fNIRS measures changes in cortical blood oxygenation and it allowed us to examine regional engagement across frontal, temporal, and parietal cortices that are relevant to higher-level language and semantic processing [12,33]. By recording these complementary electrophysiological and hemodynamic signals at the same time, we aimed to characterize how prior context shapes both rapid speech-evoked dynamics and slower, regionally distributed cortical engagement during degraded speech comprehension.

Participants listened to natural spoken narratives under three conditions: clear speech (CL), noisy speech (NS), and noisy speech preceded by prior context (NSC) (Fig. 1*A*). In the NSC condition, the prior context was provided as a brief synopsis of the upcoming story, which conveyed the global topic without directly giving answers to the subsequent comprehension questions (Fig. 1*B*). We then asked whether contextual support would modulate neural encoding at different levels of the speech-comprehension hierarchy (Fig. 1*C*). A lexical-prediction account would predict stronger or more efficient encoding of lexical surprisal and entropy when prior context is available. A semantic-scaffolding account would predict enhanced encoding of sentential semantic integration, particularly in cortical regions involved in constructing coherent discourse meaning. We also examined whether these context-sensitive neural responses were associated with comprehension performance. By combining naturalistic speech materials, simultaneous EEG-fNIRS recording, and hierarchical encoding models, this study tests how prior context shapes the neural processing of degraded speech from acoustic encoding to word-level prediction and sentence-level semantic integration. This approach allows us to ask not only whether context supports noisy speech comprehension, but also how the brain uses context to organize imperfect sensory input into meaningful communication.

**Fig. 1.**
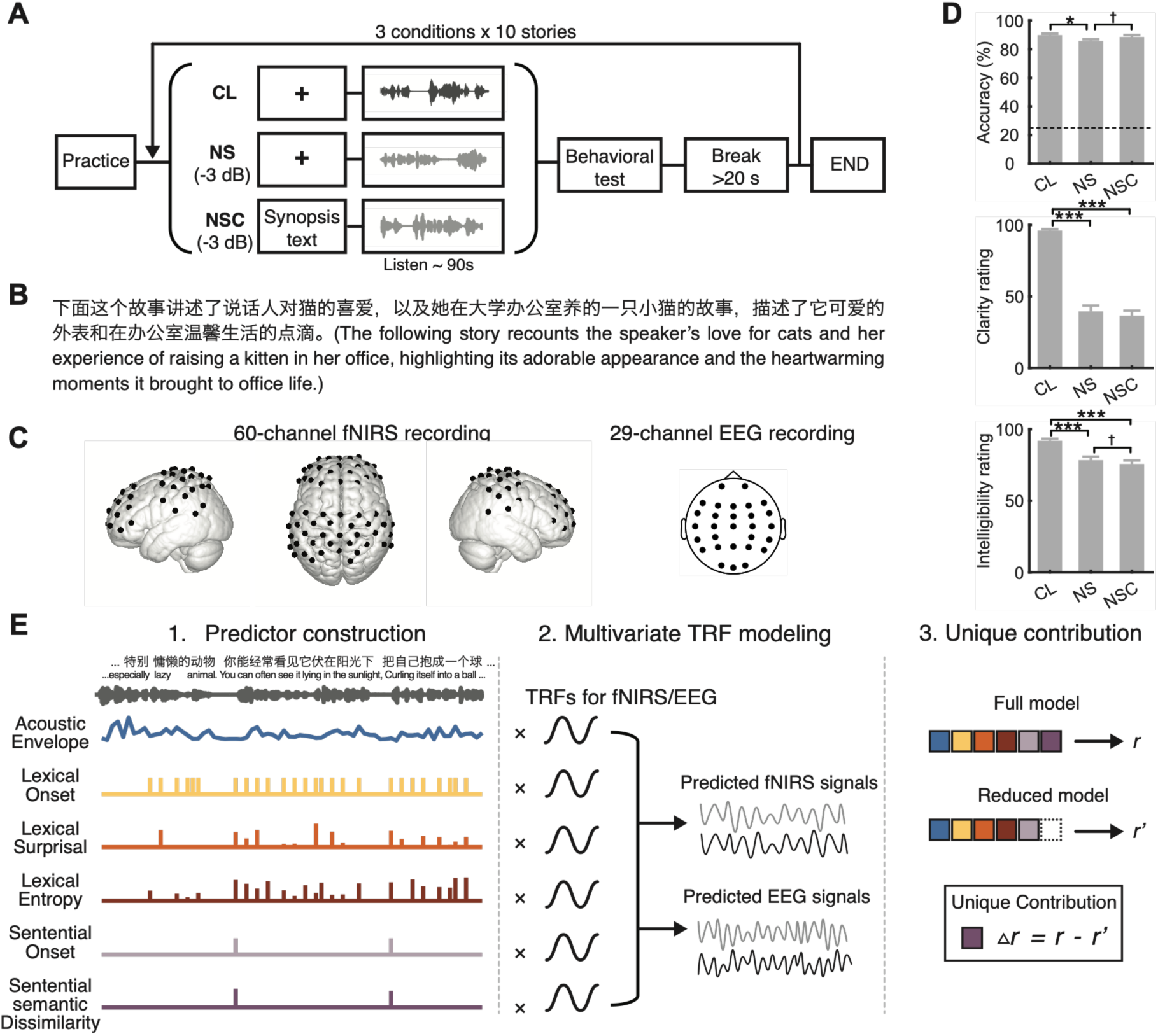
Experimental design, behavioral performance, and multivariate TRF framework. (*A*) Experimental design. Participants listened to 30 narrative recordings, each lasting approximately 90 s, under three conditions: clear speech (CL), noisy speech (NS, SNR = −3 dB), and noisy speech preceded by a prior context (NSC, SNR = −3 dB). After each narrative, participants completed a comprehension test and rated perceived clarity and intelligibility. The comprehension test included four multiple-choice questions that assessed details of the narrative content. (*B*) Example of the brief synopsis before the noisy narrative in the NSC condition. (*C*) Simultaneous EEG-fNIRS recording setup. Neural activities were recorded using 29-channel EEG and 60-channel fNIRS systems. (*D*) Behavioral comprehension accuracy and subjective clarity and intelligibility ratings across the three conditions. The dashed line indicates chance-level performance in the four-choice comprehension test. Error bars denote the standard error. (*E*) Multivariate temporal response function (mTRF) framework. Speech stimuli were represented by six predictors spanning acoustic (envelope), lexical (onset/surprisal/entropy), and sentential (onset/semantic dissimilarity) levels. These predictors were used to estimate TRFs and predict neural signals. The unique contribution of each predictor was assessed by comparing the prediction performance of the full model with that of a reduced model in which the predictor of interest was removed. †: *BF_10_* > 1, *: *BF_10_* > 3, **: *BF_10_* > 10. ***: *BF*_10_ > 100.

## Results

### Prior context facilitates noisy speech comprehension

Participants listened to 30 spoken narratives under one of three conditions: clear speech (CL), noisy speech (NS), and noisy speech preceded by prior context (NSC). In the NSC condition, prior context was provided as a brief synopsis of the upcoming story. Each narrative lasted approximately 90 s and was followed by four multiple-choice comprehension questions. Participants also rated the clarity and intelligibility of each narrative.

Speech comprehension performance was well above chance level (25%) across all conditions (CL: 89.84% ± 5.82%, mean ± SE; NS: 85.70% ± 6.79%; NSC: 88.59% ± 7.51%; Bayesian one-sample *t*-test, all *BF*_10_ > 100). Bayesian pairwise comparisons provided moderate evidence that comprehension was lower in NS than in CL (*BF*_10_ = 6.78), indicating that acoustic degradation reduced listeners’ ability to extract narrative content. Comprehension in NSC was higher than in NS, with suggestive evidence for this difference (*BF*_10_ = 2.06; Fig. 1*D*), while the comparison between CL and NSC provided moderate evidence for the absence of a difference (*BF*_10_ = 0.26; *BF_01_* = 3.85).

Subjective ratings showed a different pattern. Clarity ratings were lower in both noisy conditions than in clear speech (NS vs. CL and NSC vs. CL, both *BF*_10_ > 100), and there was modest evidence for the absence of a difference between NS and NSC (*BF*_10_ = 0.43). Intelligibility ratings were also decisively lower in both noisy conditions than in clear speech (both *BF*_10_ > 100). Moreover, there was suggestive evidence that intelligibility ratings were lower in NSC than in NS (*BF*_10_ = 2.68). These results indicate that prior context did not make the noisy speech sound clearer or more intelligible; instead, contextual support appeared to help listeners answer content-based comprehension questions despite persistently degraded subjective perceptual quality.

### Hierarchical encoding models dissociate acoustic, lexical, and sentential speech processing

To examine how prior context modulated neural encoding at different levels of speech comprehension, we used multivariate temporal response function (mTRF) models to relate continuous speech features to simultaneously recorded EEG and fNIRS signals. Six predictors were extracted from each narrative to capture acoustic, lexical, and sentential level information (Fig. 1*E*): acoustic envelope, lexical onset, lexical surprisal, lexical entropy, sentential onset, and sentential semantic dissimilarity [7,13,29,31,34]. Together, these predictors allowed us to ask whether contextual support primarily modulated low-level tracking of the speech signal, local word-level prediction, or higher-level integration of meaning across sentences.

For each predictor, we quantified its unique contribution by comparing the prediction accuracy of the full model with that of a reduced model in which the predictor of interest was removed. This approach allowed us to estimate whether a given speech feature explained neural activity beyond the variance shared with other acoustic or linguistic features. Unique contributions were summarized within predefined regions of interest (ROIs) and compared across listening conditions using Bayesian statistics.

Because EEG and fNIRS capture different types of neural signals, we used modality-appropriate summaries of the encoding results. For EEG, we examined both the unique contribution of each predictor and the temporal profile of the corresponding TRF waveform, allowing us to identify when acoustic, lexical, or sentential information was encoded. Significant temporal clusters were identified using cluster-based permutation tests, and peak amplitudes and latencies were extracted from these clusters. For fNIRS, which reflects slower hemodynamic responses, we focused on ROI-level unique contributions and used TRF waveforms primarily for visualization. This analysis framework allowed us to test whether prior context altered speech processing similarly across levels, or instead shifted the balance between lexical prediction and sentential semantic integration.

### Noise delays early acoustic-envelope encoding, with contextual modulation emerging at later stages

We first examined acoustic-envelope encoding to characterize how the brain tracked low-level amplitude fluctuations in continuous speech. EEG analysis revealed unique contributions of acoustic envelope primarily within the 0–400 ms window over central and temporal regions (Fig. 2*C*). Over central regions, cluster-based permutation tests identified an early positive TRF component at approximately 150–200 ms in all three conditions (cluster-level *p* = 0.004, 0.007, and 0.007 for CL, NS, and NSC, respectively; FDR corrected), followed by a later negative component at approximately 250–450 ms (cluster-level *p* = 0.003, 0.003, and 0.003 for CL, NS, and NSC, respectively; FDR corrected; Fig. 2*A*). Bayesian paired-sample comparisons showed decisive evidence that the latency of the early positive component was prolonged in both the NS and NSC conditions relative to the CL condition (both *BF*_10_ > 100), whereas there was moderate evidence for the absence of amplitude differences across conditions (*BF*_10_ range = 0.20–0.21). These results indicate that acoustic degradation delayed early envelope-related responses, regardless of whether prior context was available.

**Fig. 2.**
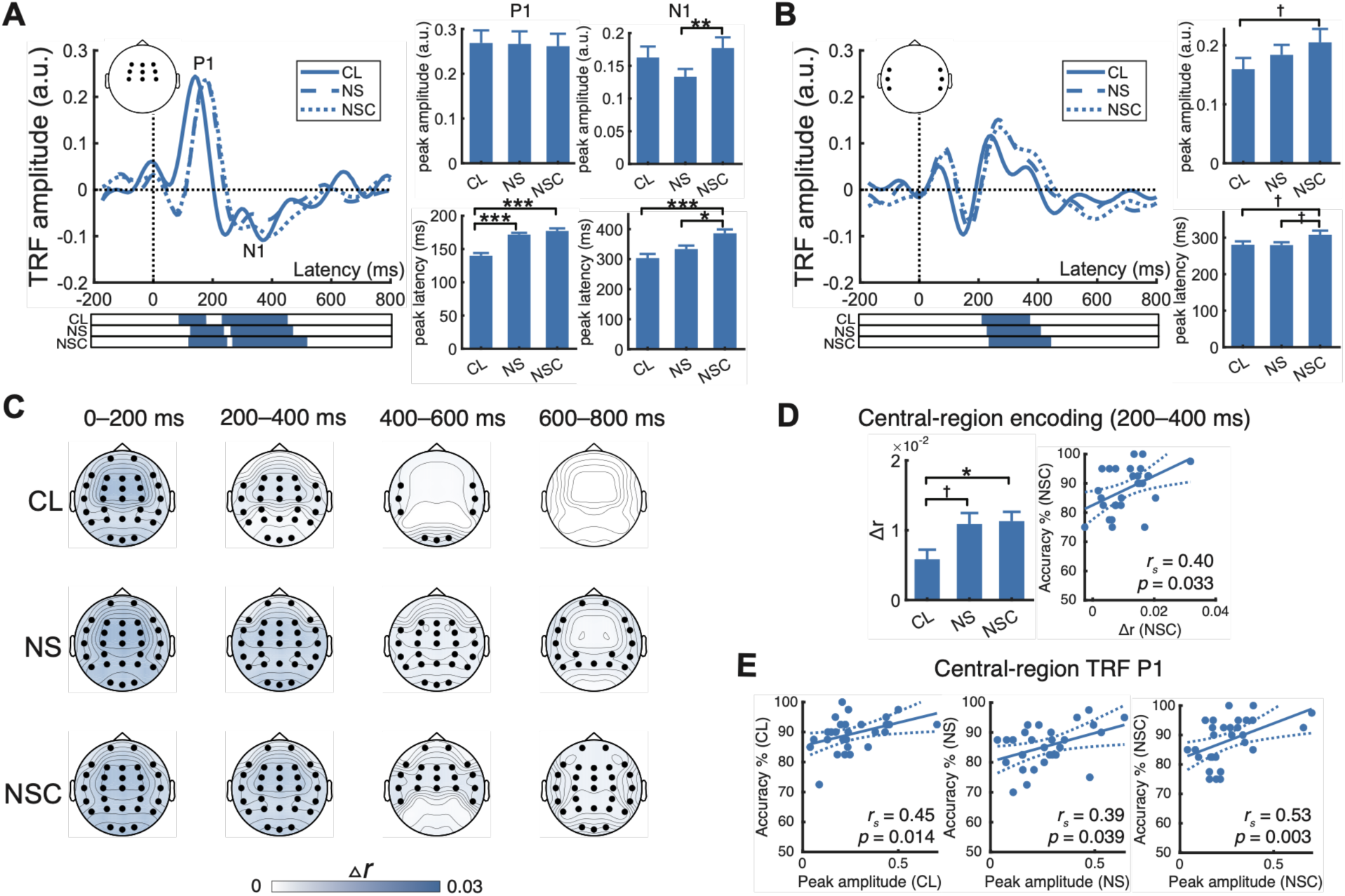
EEG-based neural encoding of acoustic envelope. (*A, B*) EEG TRFs for acoustic envelope averaged within central and temporal scalp ROIs. Solid, dashed and dotted lines indicate the CL, NS and NSC conditions, respectively. Horizontal bars below the waveforms indicate significant time windows from which peaks were extracted. Bar plots show the peak amplitude and latency. (*C*) Spatial distribution of the unique contribution of acoustic envelope across four time windows of the 0–200 ms, 200–400 ms, 400–600 ms and 600–800 ms. Black dots indicate regions showing evidence for reliable acoustic-envelope encoding. ROI-level values were projected onto the corresponding electrodes for visualization. (*D*) Unique contribution of acoustic envelope encoding in central ROI during the 200–400 ms time window and its association with behavioral comprehension accuracy in the NSC condition. (*E*) Associations between the peak amplitude of the early central acoustic-envelope TRF component and comprehension accuracy across the three listening conditions. Each dot represents one participant. Solid lines indicate linear fits and dashed lines indicate the upper and lower bounds of the confidence intervals estimated using linear regression. †: *BF_10_* > 1, *: *BF_10_* > 3, **: *BF_10_* > 10. ***: *BF*_10_ > 100.

Contextual modulation was more evident at a later stage of envelope encoding. For the later central negative component, peak latency was prolonged in NSC relative to both CL (*BF*_10_ > 100) and NS (*BF*_10_ = 6.55), and there was strong evidence for a larger peak amplitude in NSC than in NS (*BF*_10_ > 10). Thus, prior context did not restore early acoustic-envelope encoding to the clear-speech pattern, but it did modulate later envelope-related responses under noise. Over temporal regions, acoustic envelope elicited a positive TRF component at approximately 200–400 ms across all three conditions (cluster-level *p* = 0.003, 0.003, and 0.003 for CL, NS, and NSC, respectively; FDR corrected; Fig. 2*B*), with only suggestive or weak evidence for latency or amplitude differences involving NSC (latency: NSC vs. CL, *BF*_10_ = 2.22; NSC vs. NS, *BF*_10_ = 1.87; amplitude: NSC vs. CL, *BF*_10_ = 1.11).

We next examined whether acoustic-envelope encoding was associated with speech comprehension performance. In the NSC condition, stronger acoustic-envelope encoding over central region within the 200–400 ms window was associated with better behavioral comprehension accuracy (Spearman’s *r_s_* = 0.40, *p* = 0.033; Fig. 2*D*). In addition, a larger early central TRF peak was associated with higher comprehension accuracy across all three listening conditions (CL: *r_s_* = 0.45, *p* = 0.014; NS: *r_s_* = 0.39, *p* = 0.039; NSC: *r_s_* = 0.53, *p* = 0.003; Fig. 2*E*). No reliable associations were observed for peak latency, the later central component, or the temporal-region component (*p* > 0.05). These associations suggest that low-level tracking of the speech envelope remained behaviorally relevant, although the context-related effect was expressed primarily in later EEG responses rather than in a normalization of early acoustic encoding.

The fNIRS analysis provided limited evidence for acoustic-envelope encoding. Modest unique contributions were observed only in the CL condition, involving the left middle frontal gyrus (MFG) and angular gyrus (AG; *BF*_10_ = 1.71 and 1.12, respectively) and the right inferior frontal gyrus (IFG) and MFG (*BF*_10_ = 1.26 and 2.18, respectively; *SI Appendix*, Fig. S1). No ROI showed evidence for a unique contribution of acoustic envelope in the NS or NSC condition (*BF*_10_ < 0.73). Thus, the envelope results indicate that acoustic degradation affected early electrophysiological tracking of speech, while the main hemodynamic effects of prior context were not expressed at the acoustic level.

### Prior context reduces lexical prediction-related encoding under noise

We next examined whether prior context strengthened word-level predictive processing during noisy speech comprehension. Lexical surprisal quantified how unexpected each word was given the preceding context, whereas lexical entropy quantified uncertainty about likely upcoming words [7,13,29,34,35]. If contextual facilitation was primarily driven by enhanced lexical prediction, we would expect prior context to increase neural encoding of these lexical prediction-related features.

In the EEG analysis, lexical surprisal made unique contributions over temporal and parietal regions within the 400–600 ms window (Fig. 3*A*). Cluster-based permutation tests further identified an N400-like negative TRF component over parietal regions in all three conditions (cluster-level *p* = 0.003, 0.005, and 0.018 for CL, NS, and NSC, respectively; FDR corrected). Bayesian paired-sample comparisons showed that this negative component was strongly attenuated in noisy speech preceded by prior context (NSC): the peak amplitude was larger in CL than in NSC (*BF*_10_ > 100), and larger in NS than in NSC (*BF*_10_ = 3.02; Fig. 3*C*). The comparison between CL and NS provided weaker evidence for an amplitude difference (*BF*_10_ = 2.05). Thus, rather than enhancing lexical surprisal encoding under noise, prior context reduced neural sensitivity to word-level prediction error.

**Fig. 3.**
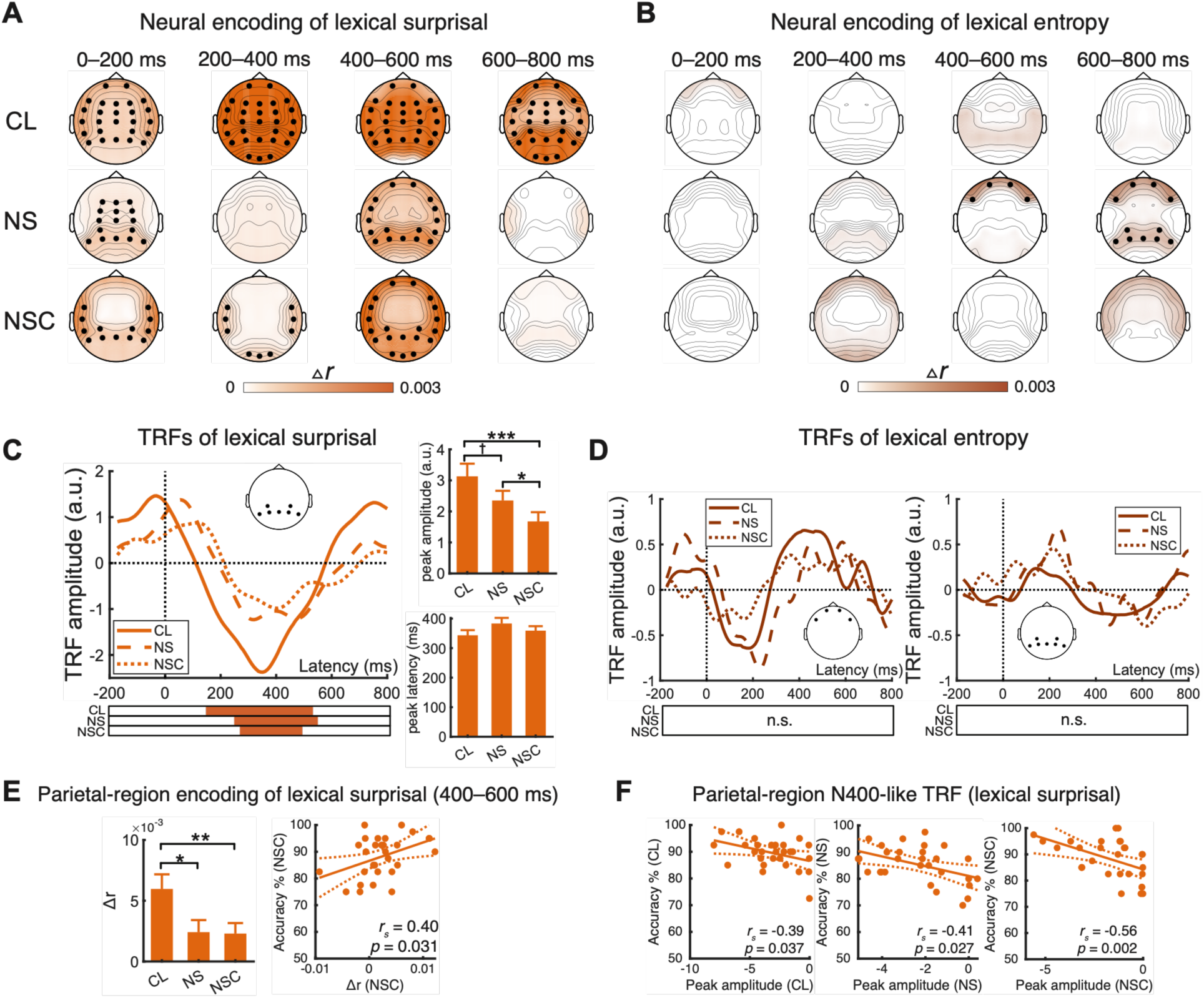
EEG-based neural encoding of lexical surprisal and lexical entropy. (*A, B*) Spatial distribution of the unique contribution of lexical surprisal (*A*) and lexical entropy (*B*) across four time windows of 0–200 ms, 200–400 ms, 400–600 ms and 600–800 ms. Black dots indicate regions showing evidence for reliable lexical surprisal/entropy encoding. ROI-level values were projected onto the corresponding electrodes for visualization. (*C, D*) EEG TRFs for lexical surprisal (*C*) and lexical entropy (*D*). Solid, dashed and dotted lines indicate the CL, NS and NSC conditions, respectively. Horizontal bars below the waveforms indicate significant time windows from which peaks were extracted. Bar plots show the peak amplitude and latency. (*E*) Unique contribution of lexical surprisal in parietal ROI during the 400–600 ms time window and its association with behavioral comprehension accuracy in the NSC condition. (*F*) Associations between the peak amplitude of the parietal lexical surprisal TRF component and comprehension accuracy across the three listening conditions. Each dot represents one participant. Solid lines indicate linear fits and dashed lines indicate the upper and lower bounds of the confidence intervals estimated using linear regression. †: *BF_10_* > 1, *: *BF_10_* > 3, **: *BF_10_* > 10. ***: *BF*_10_ > 100.

Lexical entropy made unique contributions over frontal regions within the 400–800 ms window and over parietal regions within the 600–800 ms window (Fig. 3*B*). However, no reliable TRF component was identified in either region (Fig. 3*D*). We therefore interpret the entropy results cautiously: they suggest that lexical uncertainty was represented in the EEG signal, but they do not provide strong temporal evidence that prior context enhanced uncertainty-related encoding.

We then examined whether lexical prediction-related encoding was associated with speech comprehension performance. In NSC, stronger parietal encoding of lexical surprisal within the 400–600 ms window was associated with better comprehension accuracy (Spearman’s *r_s_* = 0.40, *p* = 0.031; Fig. 3*E*). In addition, a larger parietal N400-like surprisal response was associated with higher comprehension accuracy across all three conditions (CL: *r_s_* = −0.39, *p* = 0.037; NS: *r_s_* = −0.41, *p* = 0.027; NSC: *r_s_* = −0.56, *p* = .002; Fig. 3*F*). No reliable associations were observed for peak latency (all *p* > 0.1). These associations suggest that word-level prediction-error processing remained behaviorally relevant, even though prior context reduced group-level surprisal encoding under noise.

The fNIRS results provided limited evidence for lexical prediction-related encoding. Lexical surprisal showed modest unique contributions in the right MFG in the CL (*BF*_10_ = 1.32) and in the right superior parietal gyrus (SPG) in NS (*BF*_10_ = 2.63; *SI Appendix*, Fig. S2*A*), with no ROI showing evidence for a unique contribution in NSC. Lexical entropy showed a modest unique contribution in the right IFG in NS (*BF*_10_ = 1.73), with no ROI showing evidence for a unique contribution in CL or NSC (*SI Appendix*, Fig. S2*B*). Together with the EEG findings, these results suggest that prior context did not facilitate noisy speech comprehension by strengthening lexical prediction-related encoding. Instead, contextual support was associated with attenuated tracking of word-level prediction demands, while lexical surprisal responses remained relevant for individual differences in comprehension.

### Prior context enhances fNIRS encoding of sentential semantic integration

We next examined whether prior context modulated sentential semantic integration during noisy speech comprehension. Sentential semantic dissimilarity quantified how much each incoming sentence differed from the evolving discourse context and was used to index the integration of new sentential meaning into the unfolding narrative representation. If prior context supported comprehension by providing a semantic scaffold, we expected context-supported noisy speech to show stronger encoding of this sentential semantic feature than noisy speech alone.

The EEG analysis did not reveal reliable encoding of sentential semantic dissimilarity in any condition (all *BF*_10_ < 1), suggesting that this feature was not robustly captured by electrophysiological responses in the present analysis. In contrast, the fNIRS analysis revealed condition-dependent hemodynamic encoding of sentential semantic dissimilarity across cortical regions. In CL, evidence for a unique contribution was restricted to the right IFG (*BF*_10_ = 3.17; Fig. 4*A*). In the NS condition, no ROI showed evidence for a unique contribution (all *BF*_10_ < 0.60), suggesting that acoustic degradation disrupted measurable sentential semantic encoding in the fNIRS signal.

**Fig. 4.**
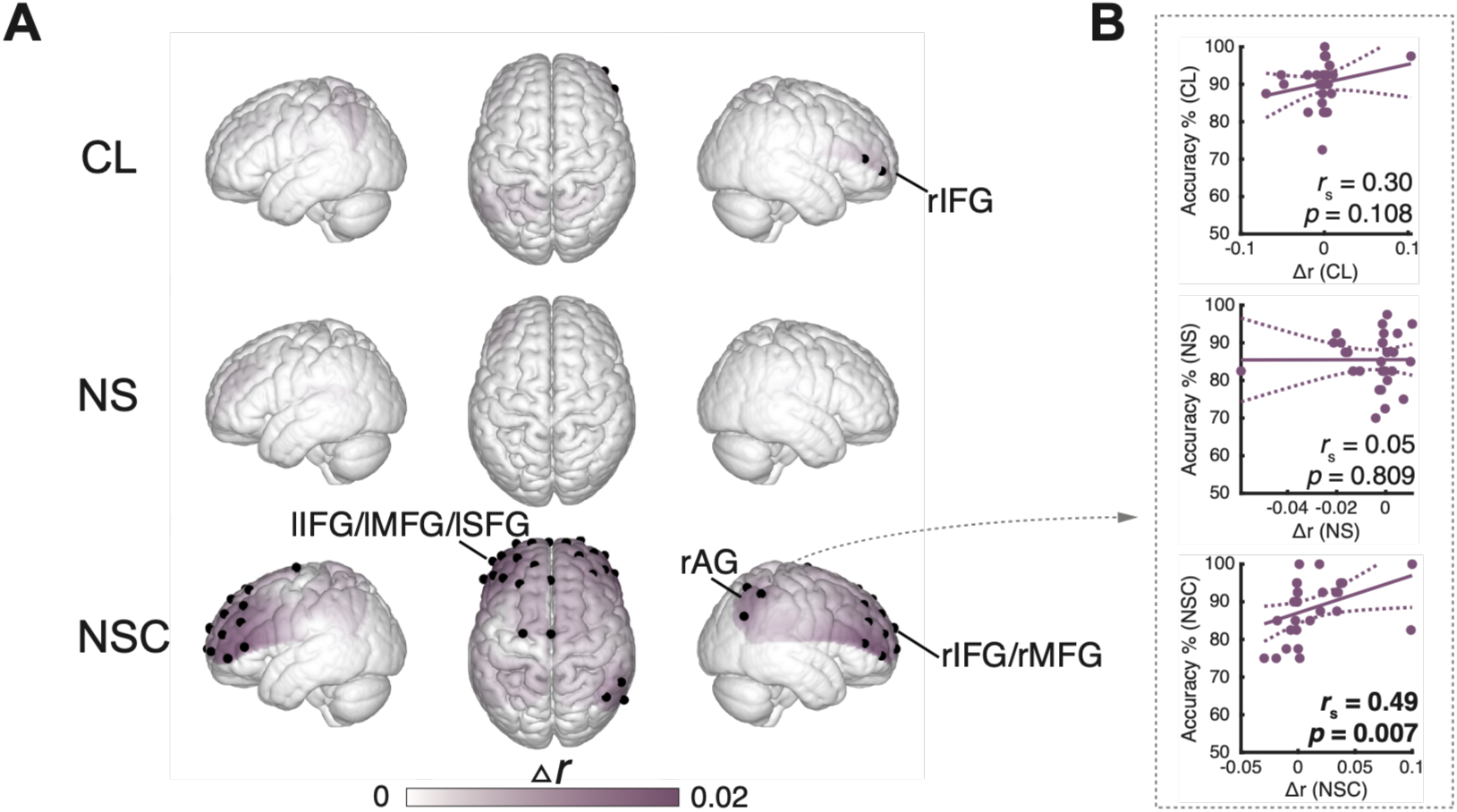
fNIRS-based neural encoding of sentential semantic dissimilarity. (*A*) fNIRS spatial distribution of the unique contribution of sentential semantic dissimilarity across the CL, NS and NSC conditions. Black dots indicate regions showing evidence for reliable sentential semantic dissimilarity encoding. ROI-level values were projected onto the corresponding channels for visualization. (*B*) Association between the unique contribution of sentential semantic dissimilarity in the rAG area and behavioral comprehension accuracy across the three listening conditions. Each dot represents one participant. Solid lines indicate linear fits and dashed lines indicate the upper and lower bounds of the confidence intervals estimated using linear regression.

When prior context was available, sentential semantic dissimilarity encoding became more broadly expressed. In NSC, evidence for a unique contribution was observed in bilateral IFG (*BF*_10_ = 3.51 and 2.84 for the left and right hemispheres, respectively), bilateral MFG (*BF*_10_ = 12.08 and 1.87), left superior frontal gyrus (SFG; *BF*_10_ = 1.50), and the right AG (*BF*_10_ = 3.10). Direct condition comparisons further revealed that right IFG encoding was lower in NS than in NSC (*BF*_10_ = 3.33), with suggestive evidence that NS was also lower than CL (*BF*_10_ = 1.79). Evidence for a difference between CL and NSC in the right IFG was weak and favored no clear difference (*BF*_10_ = 0.46; *BF*_01_ = 2.17). Together, these findings suggest that prior context was associated with a re-emergence and broadening of sentential semantic encoding under noisy listening conditions.

We then examined whether this context-sensitive sentential semantic encoding was related to comprehension performance. In the NSC condition, stronger sentential semantic dissimilarity encoding in the right AG was associated with better comprehension performance (Spearman’s *r_s_* = 0.49, *p* = 0.007; Fig. 4*B*). This association suggests that listeners who showed stronger right AG encoding of sentential semantic integration achieved better comprehension during context-supported noisy speech.

### Onset-related encoding suggests limited contextual modulation of speech boundary tracking

To determine whether the context-related effects observed for sentential semantic dissimilarity reflected broader changes in speech-boundary tracking, we further examined neural encoding of lexical and sentential onsets. Lexical onset indexed word-boundary-related processing, whereas sentential onset indexed sentence-boundary-related processing. These onset predictors allowed us to assess whether prior context generally enhanced segmentation of continuous noisy speech, or whether its effect was more specific to sentential semantic integration.

For lexical onset, EEG analysis revealed unique contributions primarily within the 200–600 ms window over central and temporal regions (*SI Appendix*, Fig. S3*C*). Over central regions, cluster-based permutation tests identified an early positive TRF component within the 0–200 ms window (cluster-level *p* = 0.028, 0.013 and 0.042 for CL, NS and NSC, respectively; FDR corrected). Bayesian comparisons showed strong evidence that the latency of this component was prolonged in NS relative to CL and shortened in NSC relative to NS (both *BF*_10_ > 10; *SI Appendix*, Fig. S3*D*), indicating that acoustic degradation delayed word-boundary-related responses while prior context mitigated this delay. Over temporal regions, lexical onset elicited a positive TRF component within the 400–600 ms window across all three conditions (cluster-level *p* = 0.031, 0.003, and 0.004 for CL, NS, and NSC, respectively; FDR corrected). This component showed moderate evidence for prolonged latency in NSC relative to CL (*BF*_10_ = 7.72), whereas the evidence for amplitude differences or latency differences between NSC and NS was weaker (NSC vs. CL, *BF*_10_ = 1.36; NSC vs. NS, *BF*_10_ = 1.77; *SI Appendix*, Fig. S3*E*). Thus, prior context selectively modulated lexical onset encoding, mainly by altering response timing, rather than producing a broad enhancement of word-boundary tracking.

The fNIRS analysis showed a complementary pattern. Lexical onset encoding was more spatially restricted in NSC than in CL or NS (*SI Appendix*, Fig. S3 *A* and *B*), suggesting that prior context did not increase widespread hemodynamic engagement for word-boundary processing. Instead, when a global semantic context was available, listeners may have required less regionally distributed engagement for segmenting the noisy speech stream into individual words. This interpretation should be treated cautiously, however, because several ROI-level effects were modest.

For sentential onset, EEG analysis revealed unique contributions between 200–600 ms over central, parietal and temporal regions across conditions (*SI Appendix*, Fig. S4*C*). However, no reliable TRF component was identified (*SI Appendix*, Fig. S4*D*). The fNIRS analysis further showed that regional involvement in sentential onset encoding was broadly similar between NS and NSC (*SI Appendix*, Fig. S4 *A* and *B*), with shared involvement of parietal and sensorimotor regions and no clear evidence that prior context generally enhanced sentential boundary encoding (NS: *BF*_10_ = 1.33, 1.21, 2.34, 2.99; NSC: 1.71, 2.52, 2.12, 3.51, for left SPG, right PoCG, SMG and SPG, respectively). No ROI showed evidence for a unique contribution of sentence onset in CL (*BF*_10_ < 0.86). These results suggest that sentence boundaries became more salient under noisy listening, but that providing prior context did not strongly amplify sentential boundary tracking.

Together, the onset-related results indicate that prior context had limited and level-specific effects on speech-boundary encoding. Lexical onset responses showed some context-related modulation, whereas sentential onset encoding was broadly similar between noisy speech with and without prior context. This pattern supports the interpretation that the broader fNIRS encoding of sentential semantic dissimilarity in NSC was unlikely to reflect a general enhancement of sentential boundary responses. Instead, prior context appeared to affect higher-level semantic integration more clearly than basic boundary tracking.

## Discussion

The present study investigated how prior context supports comprehension of noisy natural speech. By combining simultaneous EEG-fNIRS recording with hierarchical mTRF modeling, we examined how contextual support modulated neural encoding of speech information from acoustic tracking to lexical prediction and sentential semantic integration. Three main findings emerged. First, prior context brought comprehension performance under noise close to clear-speech levels, although the direct evidence for improvement over noisy speech alone was modest. Second, prior context did not enhance lexical prediction-related encoding; instead, EEG responses to lexical surprisal were attenuated in the context-supported noisy condition, even though individual differences in lexical surprisal encoding remained associated with comprehension. Third, fNIRS revealed broader encoding of sentential semantic dissimilarity across frontal regions and the right angular gyrus when prior context was available, and stronger right angular-gyrus encoding predicted better comprehension. Together, these findings suggest that prior context scaffolds noisy speech comprehension not simply by strengthening word-by-word prediction, but by helping listeners integrate degraded input into a coherent discourse-level representation.

The key advance of this study is that it identifies sentential semantic integration as a context-sensitive neural process through which prior context scaffolds comprehension of noisy natural speech. In the present study, prior context did not merely provide additional information about the upcoming story; it was associated with the re-emergence and broadening of fNIRS encoding of sentential semantic dissimilarity under noise, particularly across frontal regions and the right angular gyrus. This pattern supports the view that a global contextual framework can serve as a semantic scaffold, helping listeners organize degraded spoken input into an evolving discourse representation [14–17]. Such a scaffolding account is particularly relevant for naturalistic speech, where successful comprehension requires listeners to maintain an evolving situation model, integrate information across sentences into the broader topic of the discourse, and understand storytellers’ worldview [18,19,36,37]. The involvement of frontal regions and the angular gyrus further aligns with previous work linking these areas to discourse comprehension, internally guided semantic integration, and the use of semantic context during speech perception under adverse listening conditions [18,20,21,37–39]. Importantly, stronger right angular-gyrus encoding of sentential semantic dissimilarity predicted better comprehension in the context-supported noisy condition, suggesting that this integration-related response was behaviorally relevant. Thus, prior context may have scaffolded comprehension by helping listeners fit imperfect input into a higher-level representation of narrative meaning.

The lexical-level findings refine this scaffolding interpretation by showing that prior context did not strengthen word-by-word prediction-related encoding under noise. Lexical surprisal and entropy capture complementary aspects of predictive language processing: surprisal reflects the prediction error associated with the current word, whereas entropy reflects uncertainty about likely upcoming words [7,29,34,35]. Consistent with previous studies of naturalistic speech comprehension, lexical surprisal elicited an N400-like response over parietal regions, broadly matching the temporal profile of prediction-error processing reported in EEG and MEG studies of continuous language [7,29,35]. However, this response was attenuated when noisy speech was preceded by prior context. This pattern suggests that the synopsis did not facilitate comprehension by increasing neural sensitivity to local lexical prediction error. Rather, once listeners had access to a global semantic framework, they may have needed to rely less on continuous word-by-word prediction-error monitoring to interpret the degraded input. This interpretation is also consistent with the view that precise prediction is computationally costly and may be constrained by the need to balance prediction precision against processing efficiency [40,41]. Importantly, lexical surprisal encoding still predicted individual differences in comprehension across conditions, indicating that lexical prediction remained behaviorally relevant. Thus, the present findings do not rule out a role for predictive processing in noisy speech comprehension, especially given evidence that listeners rely more on top-down expectations when sensory evidence is unreliable [4,10,12,13]. Instead, they suggest that prior context may change how prediction and integration are balanced: local lexical prediction continues to support comprehension, but contextual facilitation in the present study was more clearly expressed as sentential semantic scaffolding.

The onset-related results further suggest that the context effect was more specific to sentential semantic integration than to general speech-boundary tracking. Lexical and sentential onsets were included as boundary-related predictors indexing the segmentation of continuous speech into word- and sentence-level units, a process that may become especially important when acoustic degradation alters the availability and reliability of segmentation cues [42–44]. In the present study, lexical onset responses showed some context-related modulation, including changes in response timing and more spatially restricted fNIRS encoding in the context-supported noisy condition. These findings are compatible with the possibility that a global semantic scaffold facilitated more efficient access to lexical boundaries while reducing the need for broadly distributed word-boundary-related processing. However, this modulation did not generalize to sentential boundary tracking: sentential onset encoding was broadly similar between noisy speech with and without prior context. This pattern contrasts with the broader fNIRS encoding of sentential semantic dissimilarity in the context-supported noisy condition, suggesting that the semantic-integration effect cannot be reduced to stronger tracking of sentence boundaries.

The distinct patterns observed in EEG and fNIRS also highlight the value of measuring complementary electrophysiological and hemodynamic signals during naturalistic speech comprehension [45,46]. In the present study, lexical surprisal effects were most clearly expressed in EEG responses, whereas context-sensitive encoding of sentential semantic dissimilarity was most clearly expressed in fNIRS responses. This dissociation is consistent with the different temporal characteristics of the processes being measured. Word-level prediction error unfolds rapidly as each word is encountered, making it well suited to EEG measures of time-sensitive electrophysiological dynamics. By contrast, sentence-level semantic integration accumulates over longer timescales and may depend on distributed cortical engagement across frontal and temporoparietal regions, making it more readily reflected in hemodynamic responses measured from the cortical areas covered by the fNIRS montage. Thus, simultaneous EEG-fNIRS recording allowed us to characterize contextual facilitation across complementary neural signals, revealing attenuated lexical prediction-related responses alongside broader cortical hemodynamic encoding of sentential semantic integration. More broadly, these findings illustrate how multimodal recording can help capture the layered nature of language comprehension, in which acoustic, lexical, and discourse-level processes unfold over different temporal scales and neural systems during real-life listening.

Several limitations should be considered when interpreting these findings. First, fNIRS is primarily sensitive to cortical regions covered by the optode montage and has limited access to deeper structures and early auditory regions, including primary auditory cortex [47,48]. Therefore, the present fNIRS results are best interpreted as reflecting hemodynamic engagement within the sampled frontal, temporal, and parietal cortices, rather than a complete account of the neural systems supporting speech-in-noise comprehension. Second, prior context was operationalized as a brief synopsis rather than as a clear version of the same passage or a verbatim transcript. This choice was intended to provide global discourse-level information while avoiding direct access to the target speech content, but different forms of prior context may engage prediction and integration mechanisms differently. Future work could compare synopsis-based, sentence-level, and word-level contextual cues to determine when prior context supports lexical prediction, semantic scaffolding, or both [4,49–51]. Finally, because lexical and sentential predictors in the context-supported condition incorporated the synopsis as preceding context, future analyses should further compare synopsis-informed and synopsis-free predictor models to clarify how global context is incorporated into computational measures of lexical uncertainty and sentential semantic integration.

In conclusion, this study shows how prior context can scaffold comprehension when speech is degraded by noise. By combining naturalistic narratives, simultaneous EEG-fNIRS recording, and hierarchical encoding models, we found that contextual support was more clearly expressed in sentence-level semantic integration than in enhanced word-by-word prediction. These findings suggest that listeners use prior context not simply to anticipate upcoming words, but to organize imperfect speech input into a coherent discourse representation. More broadly, the results highlight semantic scaffolding as a flexible neural strategy for maintaining communication under adverse listening conditions.

## Methods

### Participants

Thirty-two native Chinese speakers (17 females; mean age 23.16 years, SD = 2.23; range, 18–28 years) participated in this study. All participants were right-handed and reported normal hearing and normal or corrected-to-normal vision. The study was conducted in accordance with the Declaration of Helsinki and was approved by the Ethics Committee of Tsinghua University. Written informed consent was obtained from all participants before the experiment.

Data exclusion was performed separately for EEG and fNIRS because the two modalities had different sources of signal-quality loss. Three participants were excluded from EEG analysis because of poor EEG signal quality. fNIRS data from three additional participants were excluded, one because of poor fNIRS signal quality and two because of technical issues during recording. Thus, the EEG and fNIRS analyses each included 29 participants, with 26 participants contributing valid data for both modalities.

### Stimuli

Thirty spoken narratives were used as speech stimuli. The narratives described everyday personal experiences adapted from the National Mandarin Proficiency Test and covered 10 topic categories, such as a favorite movie or a memorable trip, with three narratives per category. They were recorded from six native Mandarin Chinese speakers with professional training in broadcasting. Recordings were made in a sound-attenuated room using a regular microphone at a sampling rate of 44,100 Hz. Each narrative lasted for around 90 s, and root mean square amplitude was matched across recordings to ensure comparable overall intensity.

For each topic category, one narrative was assigned to each listening condition: clear speech (CL), noisy speech (NS), or noisy speech preceded by prior context (NSC). This resulted in 10 narratives per condition while ensuring that topic content was balanced across conditions. The NS version was generated by adding spectrally matched stationary noise to the original clean recording at a signal-to-noise ratio of −3 dB. The noise was generated using a 50th-order linear predictive coding (LPC) model estimated from the original speech recording, following previous speech-in-noise studies [13,52].

In the NSC condition, prior context was provided as a brief written synopsis before participants listened to the noisy narrative. Each synopsis summarized the global topic and main gist of the upcoming story in approximately 50 Chinese words, without providing direct answers to the subsequent comprehension questions. The synopses were generated using ChatGPT (based on GPT4) and were manually checked to ensure that they were coherent, accurately reflected the narrative gist, and did not reveal question-specific details. An example synopsis is shown in Fig. 1B.

For each narrative, four multiple-choice questions were prepared to assess participants’ comprehension of narrative details. Each question had four response options and targeted information that required attentive listening to the narrative and could not be directly inferred from the synopsis in the NSC condition. For example, after a narrative about a speaker’s academic background, one question asked, “What is the speaker’s most likely major as a graduate student? (说话人的研究生专业最可能是什么?)” with four choices: (1) social science, (2) international politics, (3) pedagogy and (4) psychology (1. 社会科学, 2. 国际政治, 3. 教育学 and 4. 心理学).

### Experimental design

Before the formal experiment, participants completed one practice trial using an additional narrative that was not included in the main stimulus set. The practice trial was used to familiarize participants with the task procedure.

The formal experiment included 30 trials, with 10 trials assigned to each of the three conditions (CL, NS, NSC). Each narrative was presented in only one condition, and the condition assignment for each narrative was fixed across participants. Trial order was randomized separately for each participant.

In the NSC condition, a brief synopsis of the upcoming narrative was displayed on the computer screen in front of the participant before the auditory presentation. The synopsis remained visible until the participant pressed a key to proceed. During auditory presentation in all conditions, participants were instructed to maintain visual fixation to a central cross and to minimize eye blinks and all other motor activities.

After each narrative, participants answered four multiple-choice comprehension questions and rated the perceived clarity and intelligibility of the narrative on a scale from 0 to 100. Comprehension performance was quantified as the mean accuracy across trials within each condition. Participants rested for at least 20 seconds before moving to the next trial, and no performance feedback was provided.

The experimental procedure was programmed in MATLAB using the Psychophysics Toolbox 3.0 [53]. Participants listened to the narratives in a sound-attenuated room through an air-tube earphone (Etymotic ER2, Etymotic Research, Elk Grove Village, IL, USA) to reduce environmental noise and electromagnetic interference. The listening volume was adjusted to a comfortable level for each participant before the experiment and was kept consistent throughout the task. The experimental procedure is illustrated in Fig. 1 *A* and *B*.

### EEG recording and preprocessing

EEG signals were recorded simultaneously with fNIRS during the listening task using a 29-channel Neuracle EEG system (Neuracle, China). Signals were sampled at 1000 Hz. Electrodes were positioned according to the international 10–10 system, including FP1/2, FZ, F3/4, F7/8, FCZ, FC3/4, FT7/8, CZ, C3/4, T7/8, CP3/4, TP7/8, PZ, P3/4, P7/8, OZ, O1/2. Signals were recorded with CPZ as the online reference and FPZ as ground. Electrode impedances were kept below 10 kOhm for all electrodes throughout the experiment.

EEG preprocessing was performed offline in MATLAB using the FieldTrip toolbox [54]. The continuous EEG data were first notch filtered to remove 50-Hz powerline noise and re-referenced to the average of all channels. Independent Component Analysis (ICA) was then performed to identify and remove artifacts related to eye blinks and eye movements based on visual inspection. 1–6 independent components were removed per participant. The remaining components were back-projected to reconstruct the artifact-corrected EEG signals.

The preprocessed EEG signals were downsampled to 128 Hz and band-pass filtered between 1 and 8 Hz. The frequency range was selected because low-frequency EEG activity in the delta and theta bands has been widely used to examine neural tracking of continuous speech and speech comprehension under different listening conditions [30,55,56]. Finally, EEG signals were segmented into trials from 5 to 80 s relative to speech onset for subsequent encoding analyses.

### fNIRS recording and preprocessing

fNIRS signals were recorded simultaneously with EEG during the listening task using a 60-channel system (NirScan, DanyangHuichuang Medical Equipment Co., Ltd., China). Near-infrared light was emitted at three wavelengths, 730, 808, and 850 nm, and signals were sampled at 11 Hz. Optodes were positioned to cover frontal, temporal, parietal, and sensorimotor cortical regions. The Montreal Neurological Institute (MNI) coordinates and anatomical labels of all channels are provided in *SI Appendix*, Table S1.

Changes in oxy-hemoglobin (HbO) and deoxy-hemoglobin (HbR) concentration were estimated from the raw optical-density signals using the modified Beer-Lambert law. Preprocessing was performed using the Homer3 software package [57]. Motion artifacts were first corrected using a target principal component analysis algorithm (function *hmrR_MotionCorrectPCArecurse*; input parameters: tMotion = 0.5, tMask = 1, STDthresh = 30, AMPthresh = 0.5, nSV = 0.97, maxIter = 5), which identified artifact-related principal components and reconstructed the signals after removing these components [58]. To further attenuate residual motion artifacts, we then applied spline interpolation combined with Savitzky–Golay filtering (function: *hmrR_MotionCorrectSplineSG*; input parameters: p = 0.99, framesize = 10) [59]. The corrected fNIRS signals were band-pass filtered between 0.01 and 0.5 Hz (function: *hmrR_BandpassFilt*) to reduce slow drifts and high-frequency physiological noise, including heartbeat- and respiration-related fluctuations [60]. The preprocessed signals were segmented into trials from 5 to 100 s relative to speech onset.

### Predictor variables

For each narrative, six stimulus-derived predictors were extracted to represent acoustic, lexical, and sentential information (Fig. 1E). These predictors included acoustic envelope, lexical onset, lexical surprisal, lexical entropy, sentential onset, and sentential semantic dissimilarity.

At the acoustic level, the speech envelope was computed as the absolute value of the analytic signal obtained via the Hilbert transform. The resulting envelope was downsampled to 128 Hz for EEG analyses and to 11 Hz for fNIRS analyses.

At the lexical and sentential levels, each narrative was first transcribed using Iflyrec software (Iflytek Co., Ltd, Hefei, Anhui, China). Word onset times were then estimated with the Montreal Forced Aligner [61], a widely used tool for aligning speech recordings with their corresponding transcripts. These time stamps were used to construct time-aligned impulse vectors for lexical and sentential predictors.

Lexical onset was represented as a unit impulse at the onset of each word. Lexical surprisal and lexical entropy were used to quantify two complementary aspects of word-level prediction during continuous speech comprehension. Lexical surprisal indexed how unexpected the current word was given the preceding context, whereas lexical entropy indexed uncertainty about likely upcoming words [7,29,31,32,34]. Surprisal and entropy were computed as

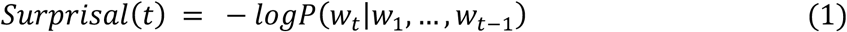

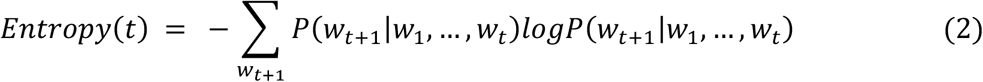

Where *w*_1_,…,*w*_(t-1)_ denotes the preceding word sequence and *P*(*w*_t_│*w*_1_,…,*w*_(t-1)_) denotes the conditional probability of the next word. To compute these values, the transcripts were stripped of punctuation and fed into a pretrained Chinese large language model (uer/gpt2-medium-chinese-cluecorpussmall) from Hugging Face [62,63], which is trained specifically to predict the next word. This model generated a probability distribution over the vocabulary for each upcoming token, from which lexical surprisal and entropy were derived [29,31,32,40].

At the sentential level, sentential onset was represented as a unit impulse at the onset of each sentence. Sentential semantic dissimilarity was used to index sentential semantic integration. Sentence embeddings were obtained using a sentence-transformer model (ibm-granite/granite-embedding-278m-multilingual) implemented in the Hugging Face environment [64]. Each sentence was represented as a 768-dimensional vector, and sentential semantic dissimilarity was defined as 1 minus the Pearson’s correlation between the vector of the current sentence and the average vector of the preceding sentence context. Higher dissimilarity values therefore indicated that the current sentence introduced semantic information that was less similar to the preceding discourse context.

Stimulus predictors were extracted from the original clean recordings for all conditions, so that neural tracking reflected the underlying speech content rather than the added noise [13,31,52]. In the NSC condition, the synopsis presented before listening was included as preceding context when computing lexical surprisal, lexical entropy, and sentential semantic dissimilarity. This allowed the linguistic predictors in the NSC condition to reflect the prior contextual information available to participants before the noisy narrative began.

The averaged values of lexical surprisal, entropy, and sentential semantic dissimilarity, along with the number of characters and sentences for each story, are reported in *SI Appendix*, Table S2. These measures did not differ significantly across conditions, indicating that the stimulus sets were broadly matched in linguistic properties (one-way ANOVA; number of words: *F*(2, 29) = 0.16, *p* = 0.854; number of sentences: *F*(2, 29) = 1.40, *p* = 0.263; lexical entropy: *F*(2, 29) = 0.73, *p* = 0.490; lexical surprisal: *F*(2, 29) = 1.37, *p* = 0.272; sentential semantic dissimilarity: *F*(2, 29) = 1.29, *p* = 0.292). Finally, lexical surprisal, lexical entropy, and sentential semantic dissimilarity were represented as impulses at their corresponding word or sentence onset times, scaled by their normalized values. Lexical onset and sentential onset were represented as unit impulses. All predictors were generated at 128 Hz for EEG analyses and at 11 Hz for fNIRS analyses.

### Stimulus-response modeling

The multivariate temporal response function (mTRF) approach based on ridge regression was employed to estimate the relationship between continuous speech features and neural responses. For each participant, condition and recording channel, stimulus predictors were used to predict the corresponding EEG or fNIRS response. Model performance was quantified as the Pearson correlation coefficient between the observed and predicted neural signals and was Fisher-*z*-transformed before statistical analysis.

The regularization parameter *λ* was selected using leave-one-trial-out cross-validation within each condition and participant. In each iteration, the model was trained on nine trials and tested on the left-out trial; this procedure was repeated until each of the ten trials in a condition had served as the test trial. Candidate *λ* values ranged logarithmically between 0.1 and 1000, and the *λ* value that yielded the highest cross-validated prediction accuracy averaged across trials was selected for subsequent modeling in that condition and participant.

For EEG analysis, TRFs were estimated over the time lags from −200 to 1000 ms, following previous studies of continuous encoding and lexical-level prediction [26,29–31]. For fNIRS analysis, TRFs were estimated over a broader time-lag window from 0 to 15 s to account for the slower hemodynamic response [60]. fNIRS trials were extended by an additional 20 s, and stimulus predictors were zero-padded accordingly to match the duration of the fNIRS response [60]. Before model fitting, neural signals were normalized to ensure consistent scaling across channels and participants, as recommended for mTRF analyses [27,28]. All mTRF analyses were implemented in MATLAB using the mTRF-Toolbox [27].

For fNIRS, the encoding analyses focused on HbO signals. HbO was selected because it generally shows a higher signal-to-noise ratio, greater sensitivity to task-related changes in cerebral blood flow than HbR, and has been used in recent studies of naturalistic speech processing, including TRF-based fNIRS work [12,60,65,66].

To quantify the unique contribution of each speech feature, we compared the prediction accuracy of a full model with that of a reduced model in which the feature of interest was excluded. The difference in prediction accuracy between the full and reduced models was taken as the unique contribution of that feature, reflecting the variance explained by the feature beyond variance shared with the other predictors [31,60]. The initial full model included all six predictors: acoustic envelope, lexical onset, lexical surprisal, lexical entropy, sentential onset, and sentential semantic dissimilarity. Predictors that did not show evidence for a unique contribution were iteratively removed, and this procedure was repeated until all predictors retained in the final model showed evidence for unique contributions. Feature-specific effects reported in the Results were based on the unique contribution of each retained predictor.

To improve signal-to-noise ratio and reduce the number of channel-wise comparisons, encoding results were summarized within predefined ROIs before statistical analysis. This ROI-level approach also allowed us to interpret context-dependent encoding patterns using predefined regional groupings rather than individual channels [66]. For EEG, channels were grouped into five scalp ROIs: frontal, central, parietal, temporal, and occipital. For fNIRS, channels were grouped into 20 anatomical ROIs based on their MNI coordinates. These ROIs included the left and right inferior frontal gyrus (IFG), middle frontal gyrus (MFG), superior frontal gyrus (SFG), precentral gyrus (PreCG), postcentral gyrus (PostCG), supramarginal gyrus (SMG), superior parietal gyrus (SPG), superior temporal gyrus (STG), middle temporal gyrus (MTG) and angular gyrus (AG).

### Statistical analyses

Statistical analyses were conducted separately for behavioral measures, EEG encoding results, and fNIRS encoding results. Bayesian one-sample or paired-sample *t*-tests were performed to quantify evidence strength using the BayesFactor MATLAB toolbox with the default Cauchy prior on effect size (scale parameter *r* = 0.707), following standard recommendations [67,68]. Evidence for the alternative hypothesis was quantified as Bayes factor *BF*_10_. Bayes factors were reported as continuous measures of evidence. Following conventional interpretive guidelines, BF values between 1 and 3 were treated as weak or suggestive evidence, values above 3 as moderate evidence, values above 10 as strong evidence and values above 100 as decisive evidence [68,69]. For behavioral analyses, comprehension accuracy, clarity ratings, and intelligibility ratings were first summarized within each condition for each participant. Bayesian one-sample *t*-tests were used to assess whether comprehension accuracy exceeded chance level. Bayesian paired-sample *t*-tests were then used to compare behavioral measures between listening conditions.

For mTRF analyses, the unique contribution of each predictor was tested at the ROI level. For each modality, condition, predictor, and ROI, Bayesian one-sample *t*-tests were used to assess whether the unique contribution was greater than zero. Bayesian paired-sample *t*-tests were used to compare predictor-specific encoding strength across conditions when relevant to the main hypotheses.

For EEG TRF waveforms, temporal clusters were identified using nonparametric cluster-based permutation tests, which provide a data-driven way to assess time-resolved effects while accounting for temporal dependence across adjacent time bins [13,29,70]. At each time bin, one-sample *t*-tests were used to examine whether the TRF amplitudes significantly differed from zero. Neighboring time bins with uncorrected *p*-value below 0.05 were grouped into clusters, and the cluster-level statistic was computed as the sum of the corresponding *t*-values. A null distribution was generated by permutations of data across participants (5,000 permutations), which defined the maximum cluster-level test statistics and corrected *p*-values for each cluster. To further control for multiple comparisons across ROIs and conditions, the resulting *p*-values were additionally corrected using the false discovery rate (FDR) procedure [71]. For significant EEG clusters, peak amplitude and peak latency were extracted for subsequent condition comparisons and brain-behavior analyses.

For fNIRS, statistical analyses focused on ROI-level unique contributions because the hemodynamic response evolves over a slower timescale and is less suited to peak-based latency analysis. fNIRS TRF waveforms were therefore used primarily for visualization (see *SI Appendix*, Fig S5), whereas statistical inference was based on cross-validated prediction accuracy and predictor-specific unique contributions.

Finally, Spearman’s rank correlation analyses were used to examine associations between neural encoding measures and comprehension performance. These brain-behavior correlations were used to assess whether context-sensitive neural responses were behaviorally relevant. Unless otherwise specified, correlation analyses were treated as exploratory and interpreted with caution.

## Data and code availability

Data and code are openly available at https://zenodo.org/records/20813685.

## Acknowledgements

This work was supported by the National Social Science Foundation of China (Major Program: 24&ZD251), the National Natural Science Foundation of China (32541018, 62577039), the Major Science and Technology Special Program of Jiangsu Province (BG2024025), Graduate Education Innovation Grants of Tsinghua University (202504Z005), and the Education Innovation Grants of Tsinghua University (DX02_20). We thank Xingda Li for help in data collection.

## Supplementary Figures

**Fig. S1.**
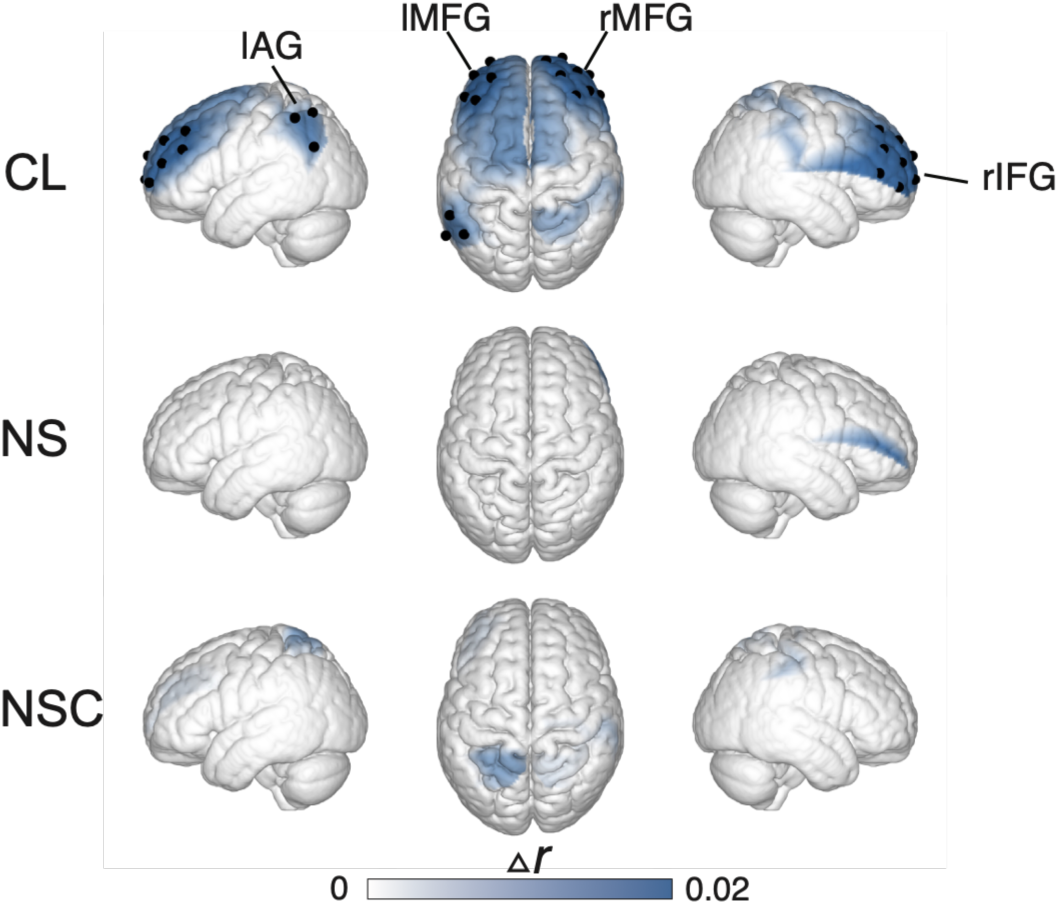
fNIRS-based neural encoding of acoustic envelope. fNIRS Spatial distribution of the unique contribution of acoustic envelope across the CL, NS and NSC conditions, indicated by black dots. ROI-level values were projected onto the corresponding channels for visualization.

**Fig. S2.**
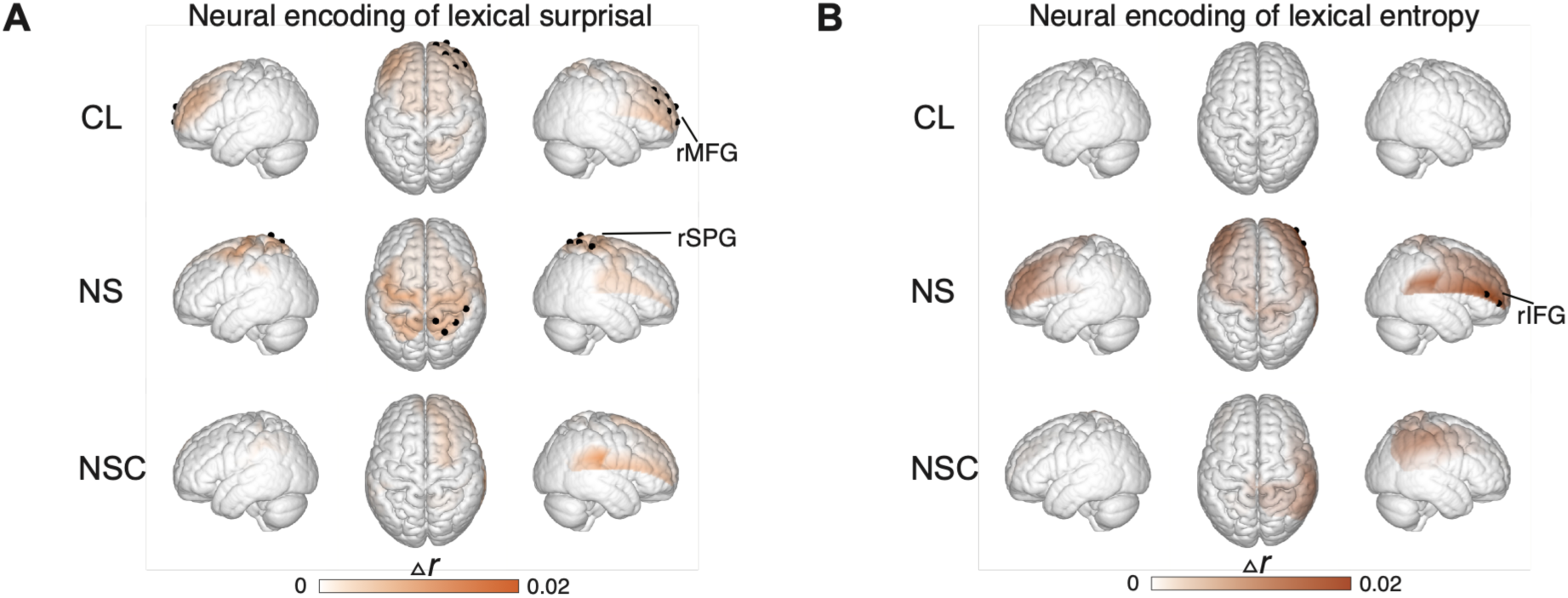
fNIRS-based neural encoding of lexical surprisal and lexical entropy. (*A, B*) fNIRS spatial distribution of the unique contribution of lexical surprisal (*A*) and lexical entropy (*B*) across the CL, NS and NSC conditions, indicated by black dots. ROI-level values were projected onto the corresponding channels for visualization.

**Fig. S3.**
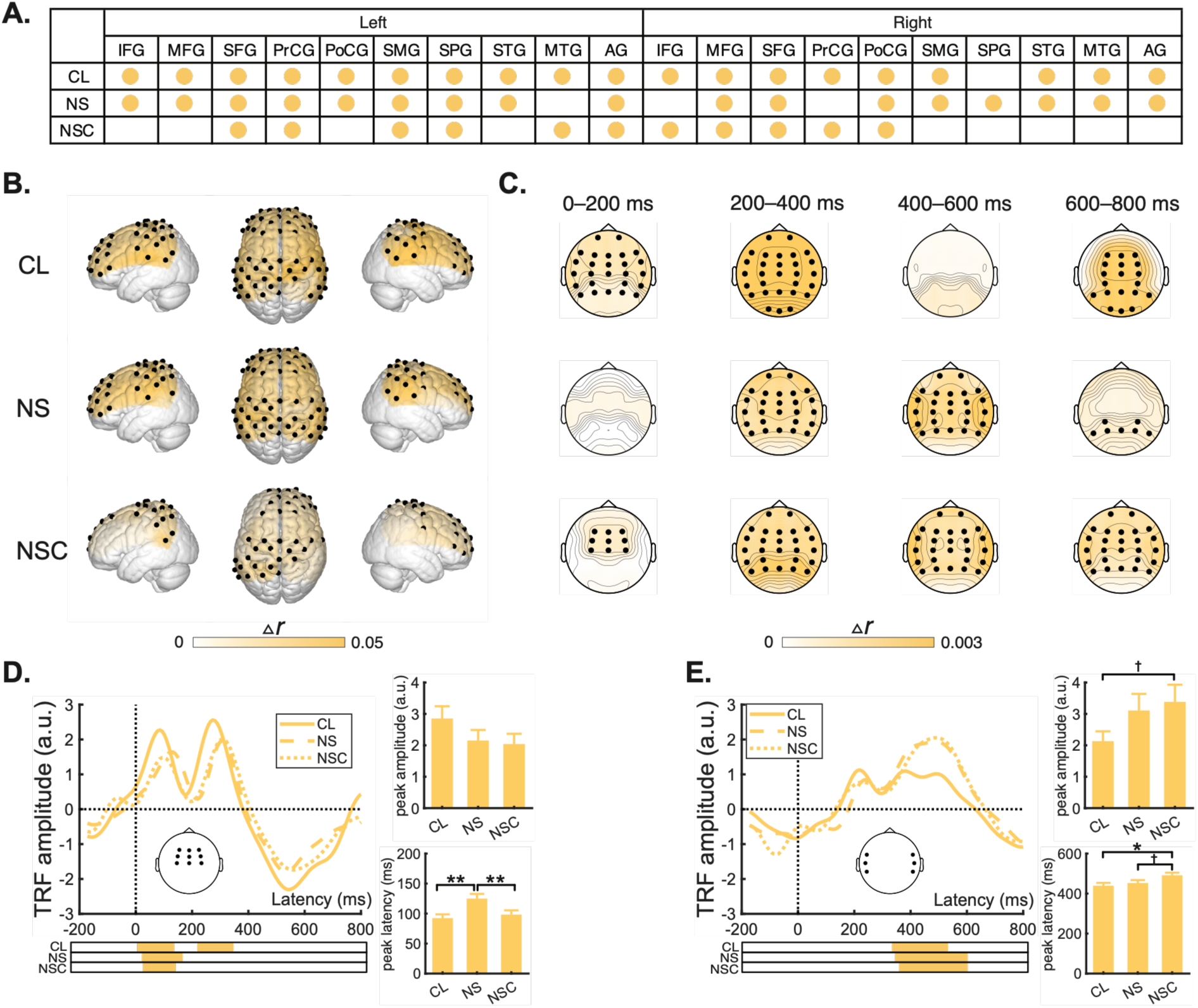
Neural encoding of lexical onset. (*A*, *B*) Summary and spatial distribution of brain regions showing unique contribution of lexical onset across the CL, NS and NSC conditions in the fNIRS analysis. These regions are indicated by colored dots in (*A*) and black dots in (*B*). ROI-level values were projected onto the corresponding channels for visualization. (*C*) EEG spatial distribution of the unique contribution of lexical onset across four time windows of 0**–**200 ms, 200**–**400 ms, 400**–**600 ms and 600**–**800 ms. Black dots indicate regions showing evidence for reliable lexical onset encoding. ROI-level values were projected onto the corresponding electrodes for visualization. (*D*, *E*) EEG TRFs for lexical onset averaged within central and temporal scalp ROIs. Solid, dashed and dotted lines indicate the CL, NS and NSC conditions, respectively. Horizontal bars below the waveforms indicate the significant time windows from which peaks were extracted. Bar plots show the peak amplitude and latency. Error bars denote the standard error. †: *BF_10_* > 1, *: *BF_10_* > 3, **: *BF_10_* > 10. ***: *BF*_10_ > 100.

**Fig. S4.**
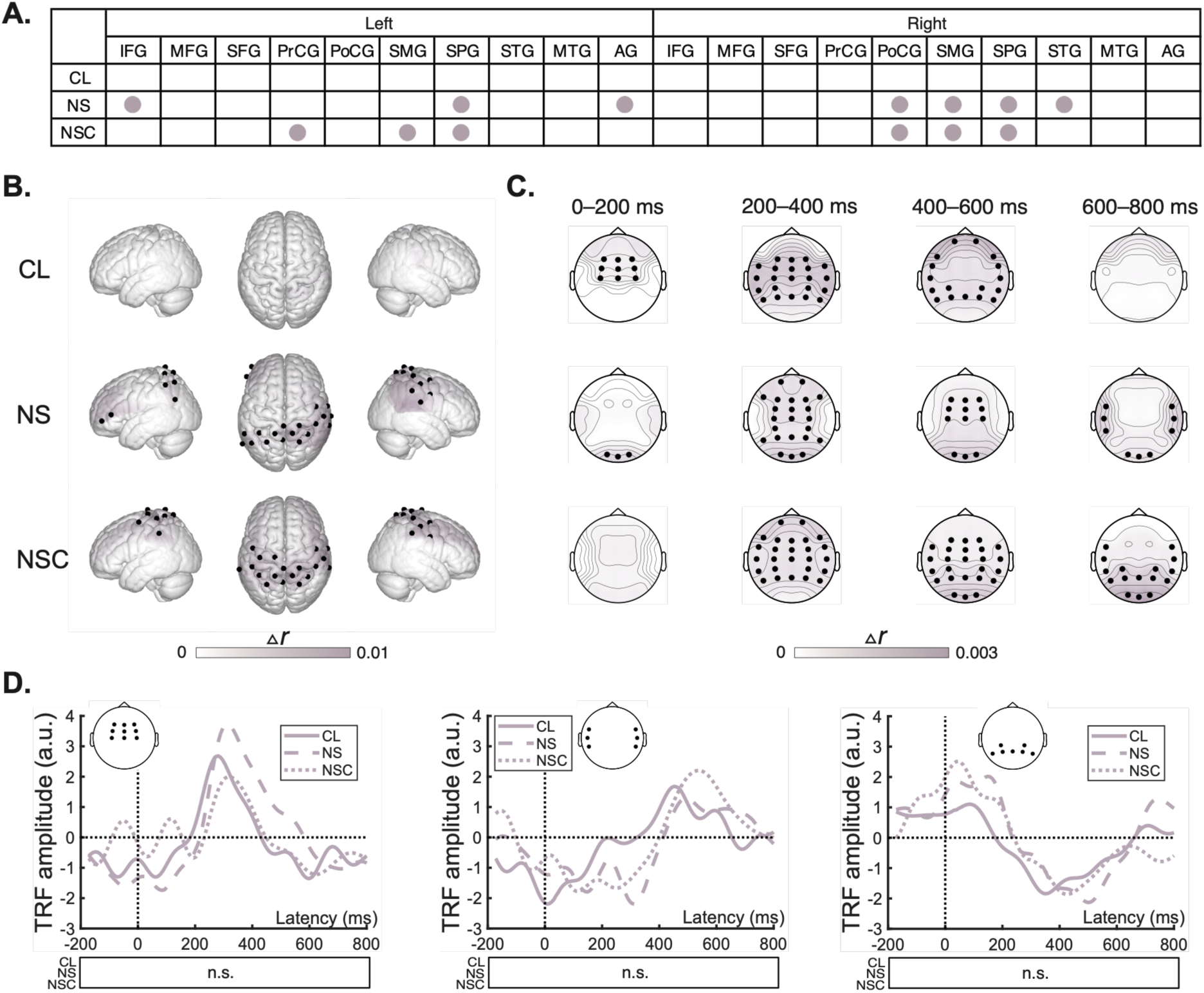
Neural encoding of sentential onset. (*A*, *B*). Summary and spatial distribution of brain regions showing unique contribution of sentential onset across the CL, NS and NSC conditions in the fNIRS analysis. These regions are indicated by colored dots in (*A*) and black dots in (*B*). ROI-level values were projected onto the corresponding channels for visualization. (*C*) EEG spatial distribution of the unique contribution of sentential onset across four time windows of 0–200 ms, 200–400 ms, 400–600 ms and 600–800 ms. Black dots indicate regions showing evidence for reliable sentential onset encoding. ROI-level values were projected onto the corresponding electrodes for visualization. (*D*) EEG TRFs for sentential onset averaged within central, temporal and parietal scalp ROIs. Solid, dashed and dotted lines indicate the CL, NS and NSC conditions, respectively. n.s. means no significant time window was identified.

**Fig. S5.**
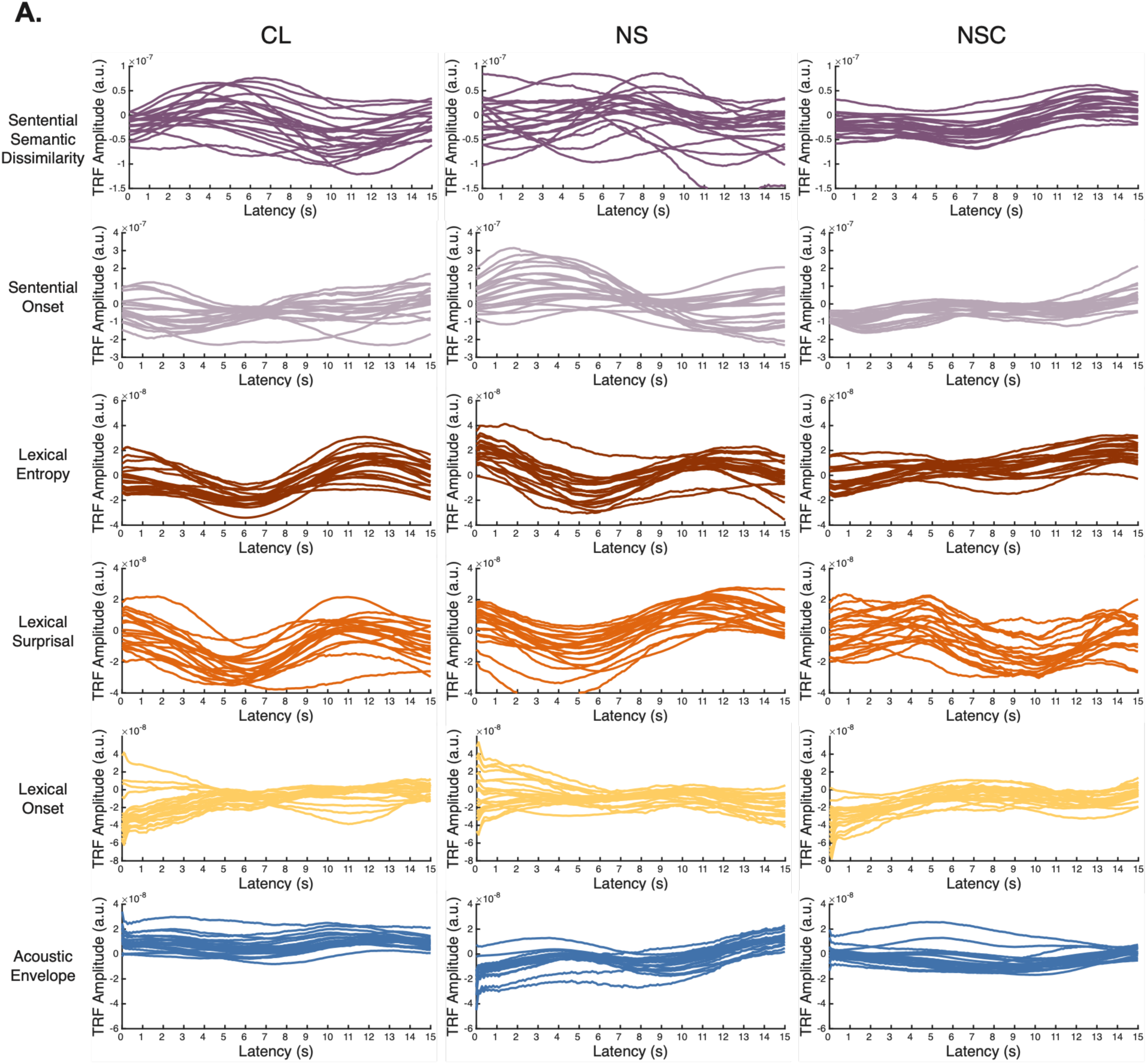
fNIRS TRFs for six predictors across the CL, NS and NSC conditions. For each predictor and condition, averaged TRFs are shown for the 20 ROIs.

## Supplementary Tables

**Table S1.**
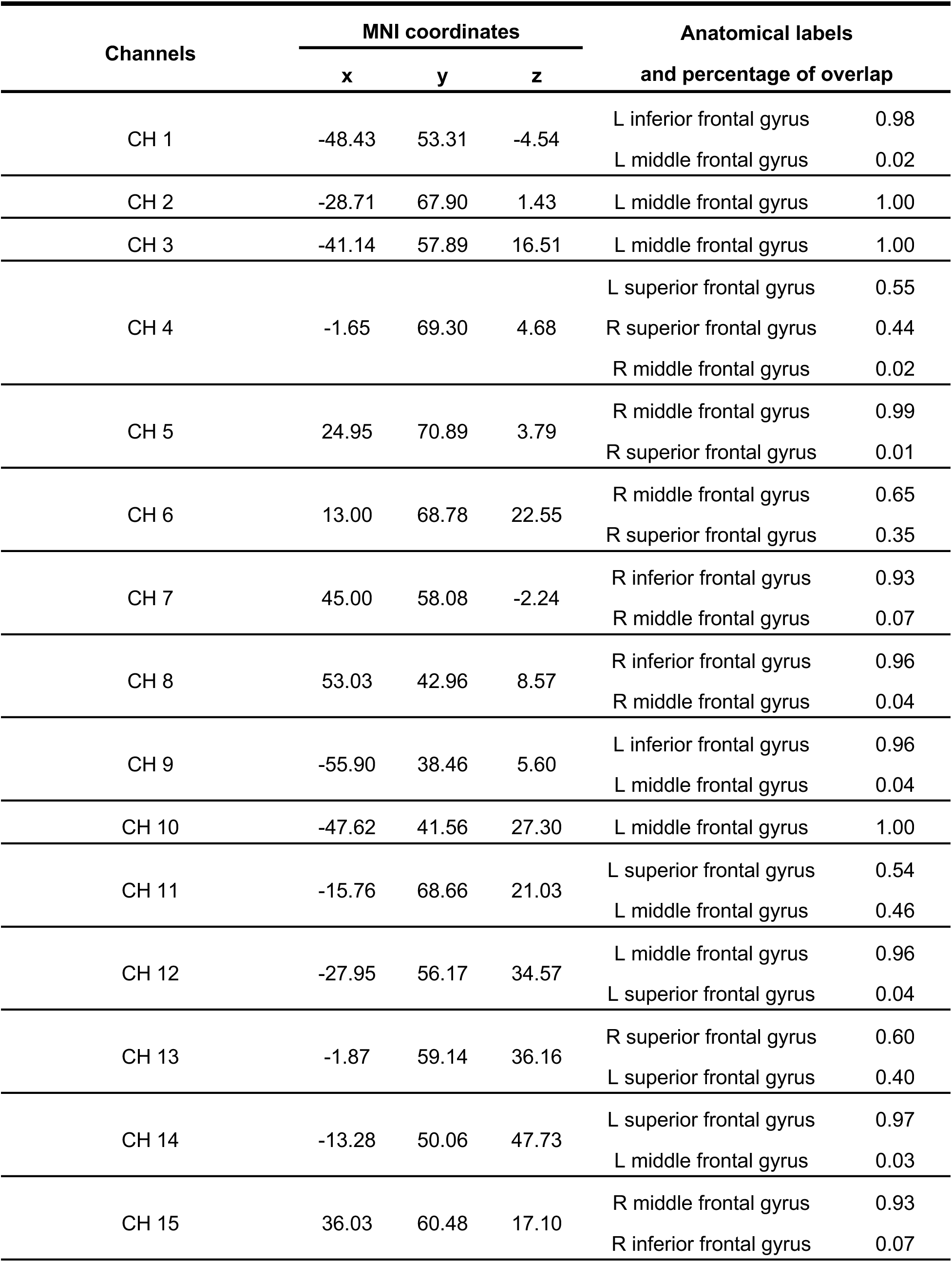
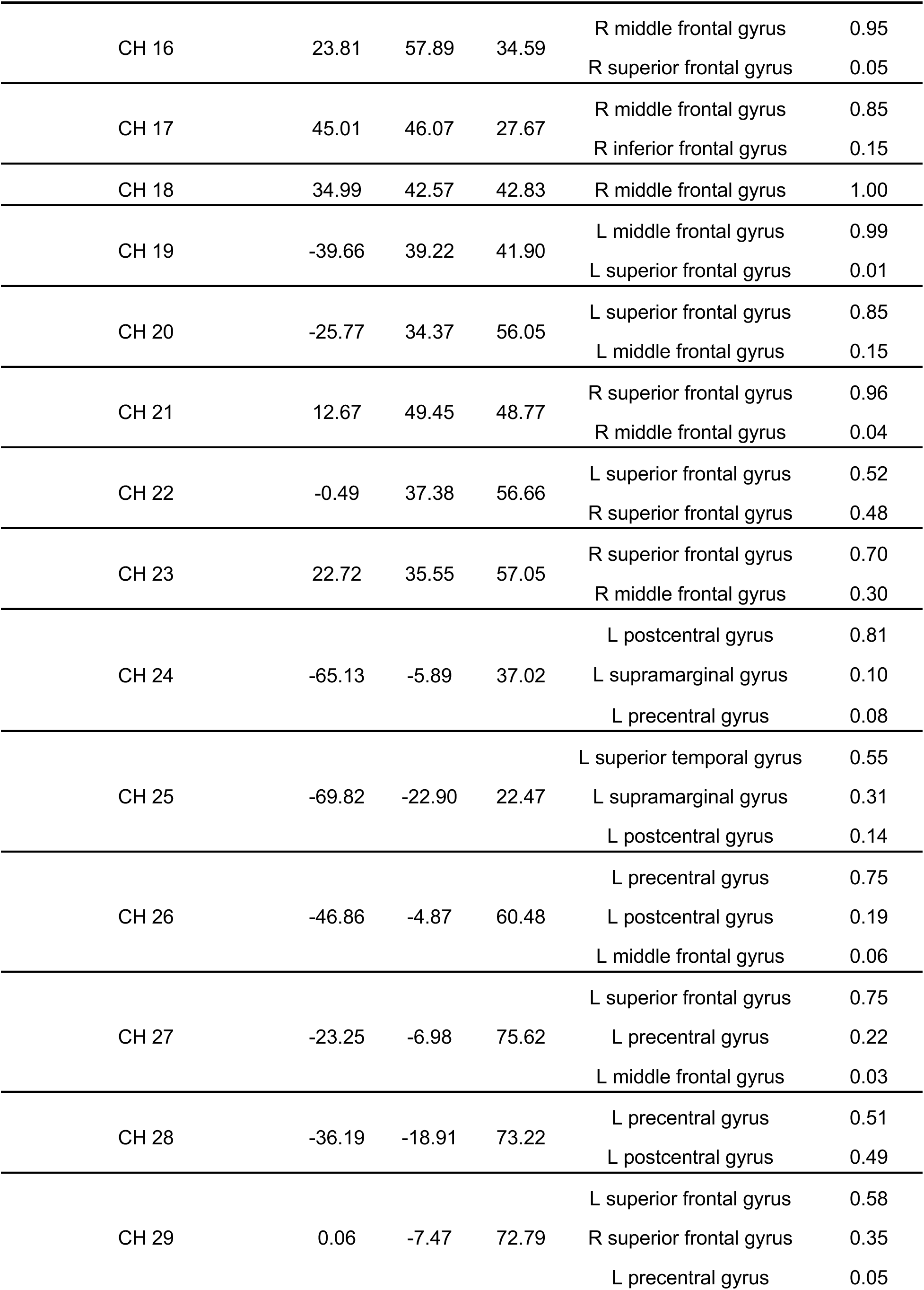
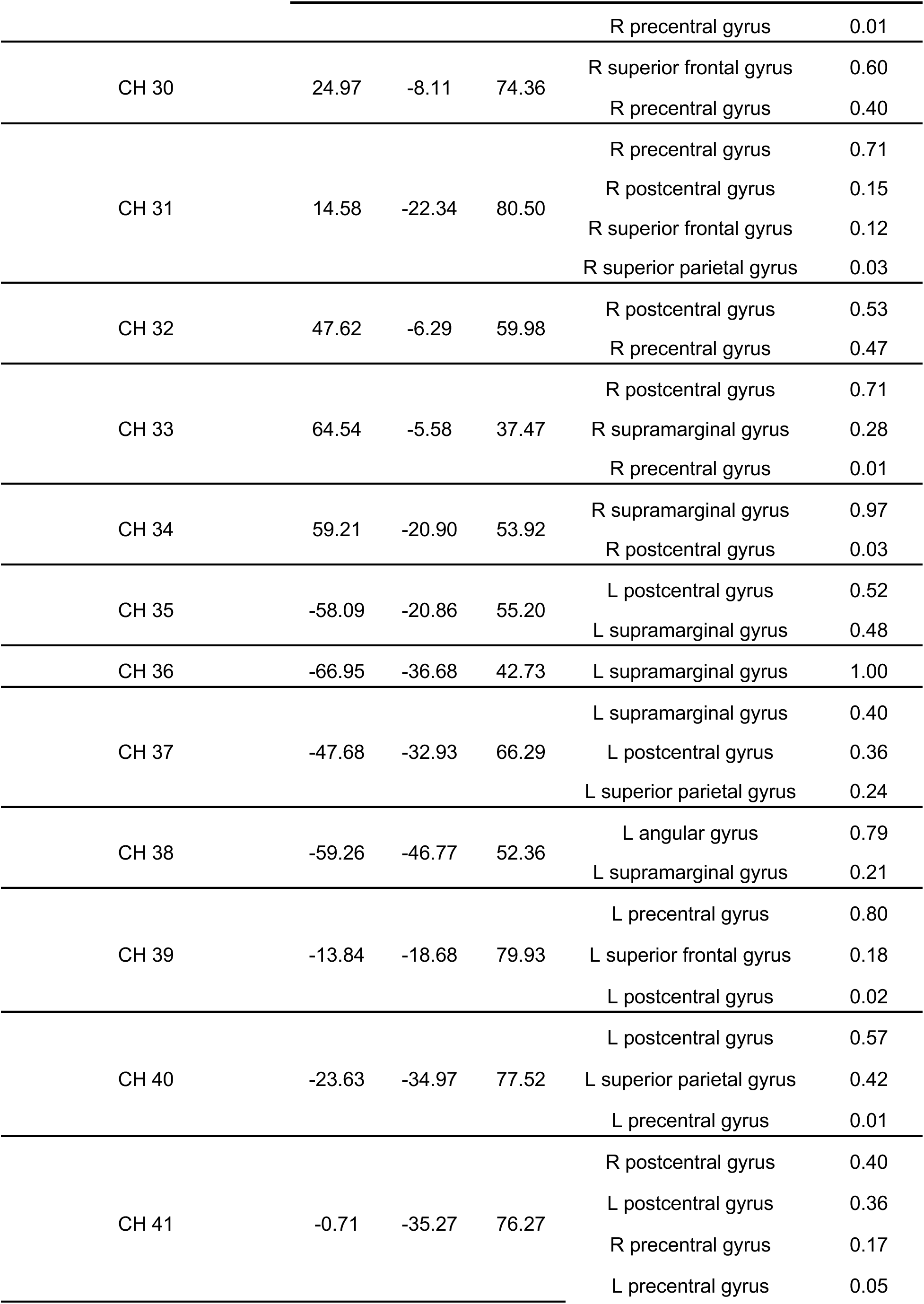
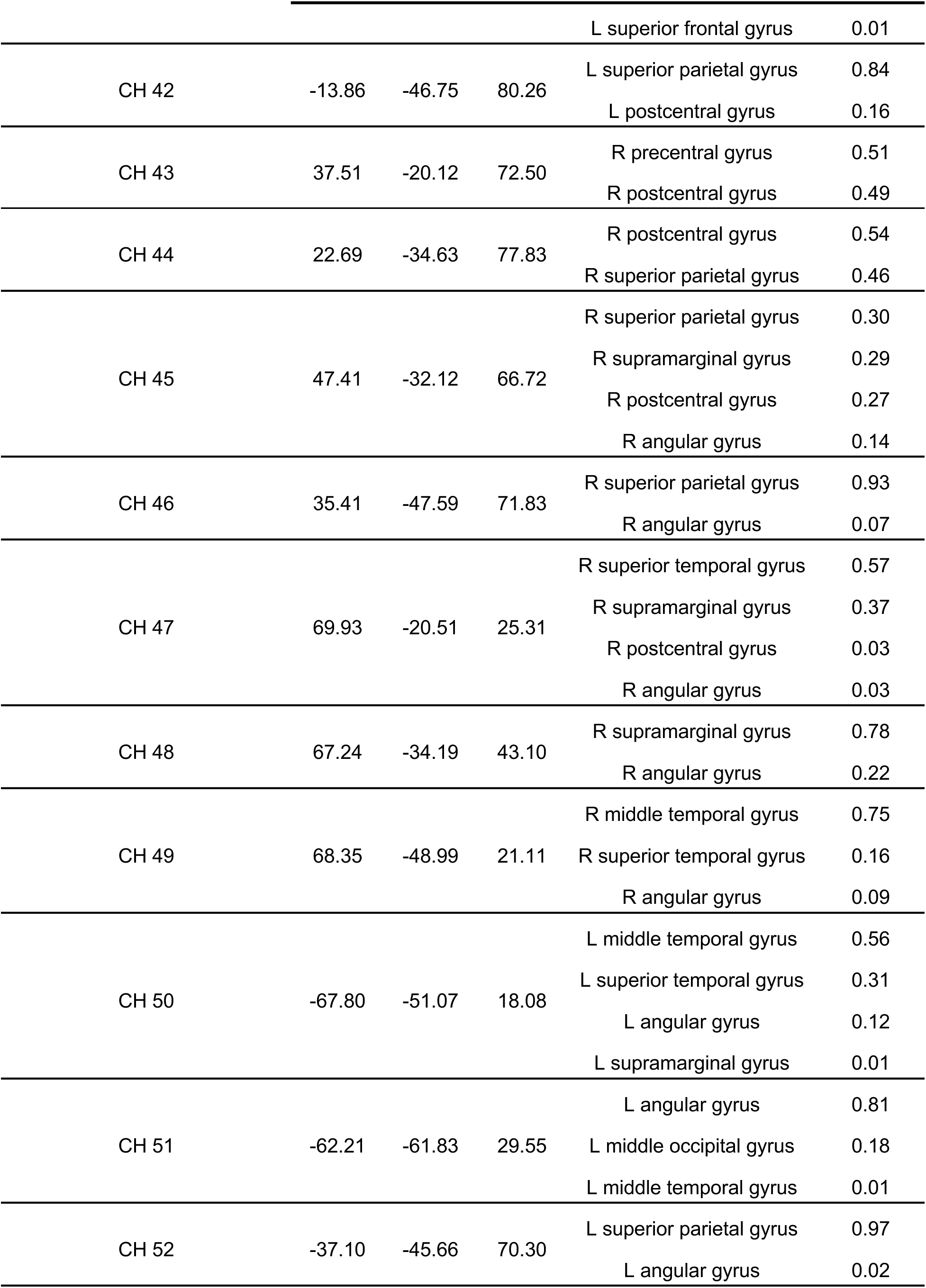
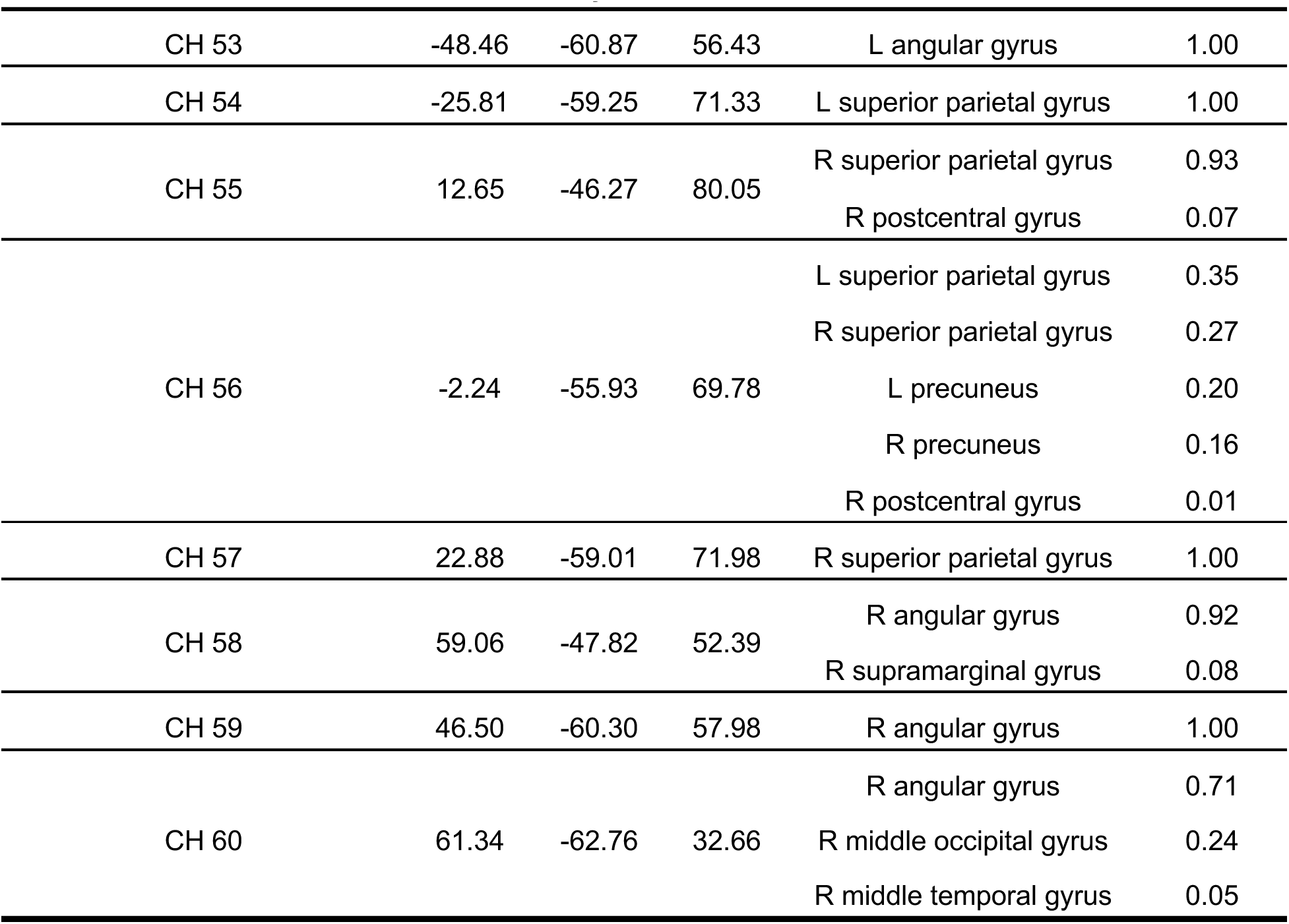
MNI coordinates and anatomical labels of fNIRS channels. The anatomical region with the maximum overlap percentage was assigned as the primary ROI label for each channel.

**Table S2.** Linguistic characteristics of the narrative stimuli across experimental conditions. Audio length, number of characters, number of sentences, averaged value and standard deviation of lexical entropy, lexical surprisal, and sentential semantic dissimilarity for each story are reported under the CL, NS, and NSC conditions.

|  | Audio<br>length (s) | Number of<br>characters | Number of<br>sentences | Lexical<br>entropy | Lexical<br>surprisal | Sentential<br>semantic<br>dissimilarity |
| --- | --- | --- | --- | --- | --- | --- |
| CL_story1 | 91.90 | 338 | 24 | 1.75(1.09) | 3.40(3.44) | 0.27(0.07) |
| CL_story2 | 91.88 | 358 | 25 | 1.74(1.06) | 3.25(3.27) | 0.26(0.07) |
| CL_story3 | 91.90 | 353 | 25 | 1.94(1.03) | 3.49(3.18) | 0.26(0.08) |
| CL_story4 | 91.88 | 384 | 26 | 1.72(1.03) | 3.08(3.18) | 0.32(0.09) |
| CL_story5 | 91.88 | 397 | 24 | 1.68(1.06) | 3.08(3.30) | 0.28(0.08) |
| CL_story6 | 91.90 | 414 | 29 | 1.86(1.04) | 3.60(3.56) | 0.27(0.07) |
| CL_story7 | 91.90 | 364 | 31 | 1.97(1.02) | 3.94(3.56) | 0.26(0.07) |
| CL_story8 | 91.88 | 386 | 27 | 1.66(1.10) | 3.39(3.62) | 0.29(0.08) |
| CL_story9 | 91.88 | 429 | 36 | 1.90(1.06) | 3.43(3.25) | 0.27(0.07) |
| CL_story10 | 92.00 | 378 | 27 | 1.69(1.14) | 3.22(3.47) | 0.25(0.06) |
| NS_story1 | 91.90 | 366 | 33 | 1.66(1.02) | 3.03(3.30) | 0.25(0.07) |
| NS_story2 | 91.90 | 380 | 25 | 1.79(1.07) | 3.65(3.63) | 0.27(0.08) |
| NS_story3 | 91.88 | 416 | 31 | 1.82(1.07) | 3.52(3.47) | 0.27(0.08) |
| NS_story4 | 91.88 | 356 | 30 | 1.79(1.10) | 3.33(3.29) | 0.27(0.07) |
| NS_story5 | 91.90 | 364 | 29 | 1.74(1.11) | 3.34(3.59) | 0.27(0.07) |
| NS_story6 | 91.88 | 353 | 29 | 1.74(1.09) | 3.23(3.35) | 0.25(0.07) |
| NS_story7 | 91.90 | 348 | 29 | 1.81(1.01) | 3.60(3.52) | 0.26(0.07) |
| NS_story8 | 91.88 | 427 | 34 | 1.63(1.08) | 3.29(3.61) | 0.27(0.08) |
| NS_story9 | 91.88 | 485 | 34 | 1.77(1.08) | 3.56(3.55) | 0.26(0.07) |
| NS_story10 | 91.88 | 364 | 27 | 1.78(1.03) | 3.26(3.23) | 0.27(0.08) |
| NSC_story1 | 91.88 | 367 | 33 | 1.85(1.04) | 3.57(3.41) | 0.28(0.08) |
| NSC_story2 | 91.88 | 365 | 31 | 1.82(1.02) | 3.39(3.39) | 0.28(0.08) |
| NSC_story3 | 91.88 | 388 | 26 | 1.72(1.04) | 3.60(3.76) | 0.27(0.09) |
| NSC_story4 | 91.88 | 431 | 34 | 1.85(1.09) | 3.76(3.58) | 0.28(0.07) |
| NSC_story5 | 91.88 | 398 | 25 | 1.76(1.03) | 3.44(3.60) | 0.28(0.08) |
| NSC_story6 | 91.88 | 398 | 29 | 1.90(1.05) | 3.67(3.47) | 0.25(0.07) |
| NSC_story7 | 91.88 | 328 | 24 | 1.87(1.05) | 3.79(3.61) | 0.27(0.08) |
| NSC_story8 | 91.88 | 435 | 33 | 1.79(1.09) | 3.77(3.85) | 0.28(0.07) |
| NSC_story9 | 91.88 | 396 | 25 | 1.86(1.06) | 3.47(3.36) | 0.25(0.09) |
| NSC_story10 | 91.90 | 382 | 23 | 1.69(1.11) | 3.16(3.46) | 0.26(0.08) |

